# Fluorescence exclusion: a rapid, accurate and powerful method for measuring yeast cell volume

**DOI:** 10.1101/2021.10.07.463508

**Authors:** Daniel García-Ruano, Akanksha Jain, Joseph C. Ryan, Vasanthakrishnan Radhakrishnan Balasubramaniam, Larisa Venkova, Matthieu Piel, Damien Coudreuse

## Abstract

Cells exist in an astonishing range of volumes across and within species. However, our understanding of cell size control remains limited, due in large part to the challenges associated with accurate determination of cell volume. Much of our comprehension of size regulation derives from models such as budding and fission yeast, but even for these morphologically stereotypical cells, assessment of cell volume has relied on proxies and extrapolations from two-dimensional measurements. Recently, the fluorescence exclusion method (FXm) was developed to evaluate the size of mammalian cells, but whether it could be applied to smaller cells remained unknown. Using specifically designed microfluidic chips and an improved data analysis pipeline, we show here that FXm reliably detects subtle difference in the volume of fission yeast cells, even for those with altered shapes. Moreover, it allows for the monitoring of dynamic volume changes at the single-cell level with high time resolution. Collectively, our work reveals how coupling FXm with yeast genetics will bring new insights into the complex biology of cell growth.

**SUMMARY STATEMENT:** Fluorescence exclusion provides a unique method to accurately measure the volume of yeast cells at both the population and single-cell levels.

## INTRODUCTION

Cell volume is a dynamic and complex trait that is modulated by both genetic and environmental components (Amodeo and Skotheim, 2016; Cook and Tyers, 2007; Lloyd, 2013; Marshall et al., 2012; Mueller, 2015). It has an impact on a host of cellular processes (Pedersen et al., 2001; Zhurinsky et al., 2010), playing important roles in tissue architecture and contributing to the overall size and shape of an organism (Amodeo and Skotheim, 2016; Cook and Tyers, 2007; Lloyd, 2013; Marshall et al., 2012; Mueller, 2015). At the single-cell level, volume and surface area not only define how cells sense external stimuli and interact with their environment but also determine their intracellular chemistry and organization. Strikingly, deregulation of cell size has dramatic consequences and is a hallmark of aging and cancer (Li et al., 2015; Pietras, 2011; Yang et al., 2011).

In proliferating cells, volume is intimately linked to the control of cell cycle progression. The dynamics of cell growth are complex across species, and specific stages of the cell cycle are associated with differential changes in cell size and morphology. For instance, while cells of the fission yeast *Schizosaccharomyces pombe* stop growing and maintain their overall shape during mitosis, mammalian cells undergoing division show a transient phase of rounding and volume alteration (Lancaster et al., 2013; Zlotek-Zlotkiewicz et al., 2015). In addition, size thresholds have been proposed to act as key checkpoints during the fission yeast cell cycle, allowing for the coordination of growth and division (Fantes, 1977; Fantes and Nurse, 1978; Fantes and Nurse, 1977). These mechanisms reduce cell-to-cell heterogeneity, thereby maintaining size homeostasis in a growing population.

Alterations in cell size are also triggered by changes in the environment, such as upon modulation of nutritional conditions, and cell cycle exit is often associated with a decrease in cell volume. For example, yeast cells exposed to nitrogen or glucose starvation enter quiescence with a significantly reduced size (Sun and Gresham, 2021). Interestingly, increases in cell volume were frequently observed during long-term experimental evolution in bacteria and were proposed to contribute to improved proliferation (Grant et al., 2021; Lenski and Travisano, 1994; Mongold and Lenski, 1996). This highlights how the volume of dividing cells may result from the necessary balance between growth advantages and evolutionary trade-offs.

The principles and mechanisms underlying cell size control have been the focus of in-depth investigation. While several pathways have been described to play a role in regulating this critical parameter (Amodeo and Skotheim, 2016; Cook and Tyers, 2007; Lloyd, 2013; Marshall et al., 2012; Mueller, 2015), whether they represent the key mechanisms that ensure size homeostasis remains unclear. Furthermore, how cells measure and control their geometric characteristics is surprisingly poorly understood. As a result, multiple models that are not mutually exclusive have been proposed: cells may divide 1) when they have reached a specific size (sizer), 2) after a specific time following a landmark cell cycle event (timer) and 3) after having produced a fixed amount of mass, at a rate that depends on birth size (adder). While these possibilities are supported by experimental and theoretical studies (Fantes and Nurse, 1977; Mueller, 2015; Soifer et al., 2016; Sveiczer et al., 1996; Taheri-Araghi et al., 2015), general conclusions are difficult to draw. This may be due in part to the diversity of measurements used to describe cell size, including volume, surface area, cell length or dry mass. Indeed, the question remains whether one or a combination of these characteristics is more relevant for understanding size regulation and whether different cell types rely on the same or on distinct parameters. Thus, while the sizes and shapes of individual cell types are commonly accepted as fundamental hallmarks of cell identity, no unifying principles for size control have emerged.

Experimentally, the regulation of cell size has been difficult to investigate. In mammalian cell lines, the challenges of advanced genetic manipulation and the lack of simple methods for accurate measurement of parameters such as volume and surface area have been an obstacle to studying the molecular mechanisms of size control. Therefore, unicellular eukaryotes such as the budding and fission yeasts remain the models of choice for deciphering the bases of this critical cellular feature. In these organisms, regulation of cell size is intimately linked to the operation of the cell cycle machinery. Different concepts have thus emerged that may provide robust ways of regulating size, including titration mechanisms and dependence on geometric parameters (*e.g*. length, surface area) (Amodeo and Skotheim, 2016; Facchetti et al., 2019). However, how cells monitor their dimensions and how variation from the target size triggers a corrective response are not fully understood.

In the fission yeast *S. pombe*, the existence of cell size checkpoints was initially suggested at both S phase and, more importantly, mitosis (Fantes, 1977; Fantes and Nurse, 1978; Fantes and Nurse, 1977). While a mechanism based on an intracellular gradient of a cell cycle regulator was subsequently proposed (Martin and Berthelot-Grosjean, 2009; Moseley et al., 2009), recent evidence has challenged this model (Amodeo and Skotheim, 2016; Pan et al., 2014a; Saunders et al., 2012). Moreover, the commonly-employed approach of using cell length at division as a proxy for size may not be sufficient for investigating size regulation. First, cell volume is more critical to cell biology and biochemistry than length. Second, slight changes in the diameter of these rod-shaped cells are usually ignored, although they have a stronger impact on cell volume than similar alterations in length: the volume of a cylinder is linearly dependent on its length but varies with the square of its radius. In addition, morphology mutants or cells grown in conditions that induce alteration of cellular dimensions and scaling are complex to study and to compare with reference strains using this method. Finally, size at division restricts the analysis to only a small fraction of a proliferating population.

To investigate the control of yeast cell volume in a more unbiased manner, different techniques have been applied. These include volume calculation from geometrical assumptions, 2D imaging data that allow for 3D reconstruction or the integration of cellular sections extrapolated from the cell outlines, and complex approaches using advanced micromechanical devices (Baybay et al., 2020; Bryan et al., 2010; Bryan et al., 2014; Facchetti et al., 2019; Model, 2018; Pan et al., 2014b; Zegman et al., 2015). However, these strategies are either low-throughput, difficult to establish or rely on various assumptions and complex processing of 2D measurements. For instance, given the impact of cell diameter on cell volume, accurate evaluation of cell size based on automated recognition of cell outlines requires this detection to be highly precise (Baybay et al., 2020; Zegman et al., 2015). Therefore, while these methods represent important steps in our capacity to address the regulation of cell volume, a more direct and simple methodology that does not introduce the biases of 2D data analysis or multiple steps of image segmentation and reconstruction is necessary.

Recently, a fluorescence exclusion method (FXm) was developed for evaluating the volume of morphologically diverse mammalian cells (Cadart et al., 2017). FXm relies on microfluidic devices in which cells are imaged in the presence of a fluorescently-labelled dextran that does not cross the plasma membrane. Thus, using chambers of known height, the local reduction in fluorescence due to the presence of a cell allows for the determination of the volume of this cell with high accuracy (Fig. 1A). This technique is simple and only requires single-plane imaging with low magnification objectives. It is unaffected by morphological alterations and is compatible with the monitoring of high numbers of cells. Although perfectly adapted to large mammalian cells, whether this new approach could be used for smaller cells such as yeast remained unclear, as this would require microdevices with significantly lower internal chambers. For example, HeLa cells are around 20-40 μm in height with a volume of ~1500 μm^3^ at birth and ~3000 μm^3^ at division (Cadart et al., 2018; Cadart et al., 2019). In contrast, wild-type fission yeast cells divide at a length of ~14 μm, with a diameter of ~4-4.5 μm. Depending on the growth conditions and exact diameter used for volume calculations, this led to values for size at division between ~120 and 200 μm^3^ (Navarro and Nurse, 2012; Nurse, 1975).

**Figure 1.**
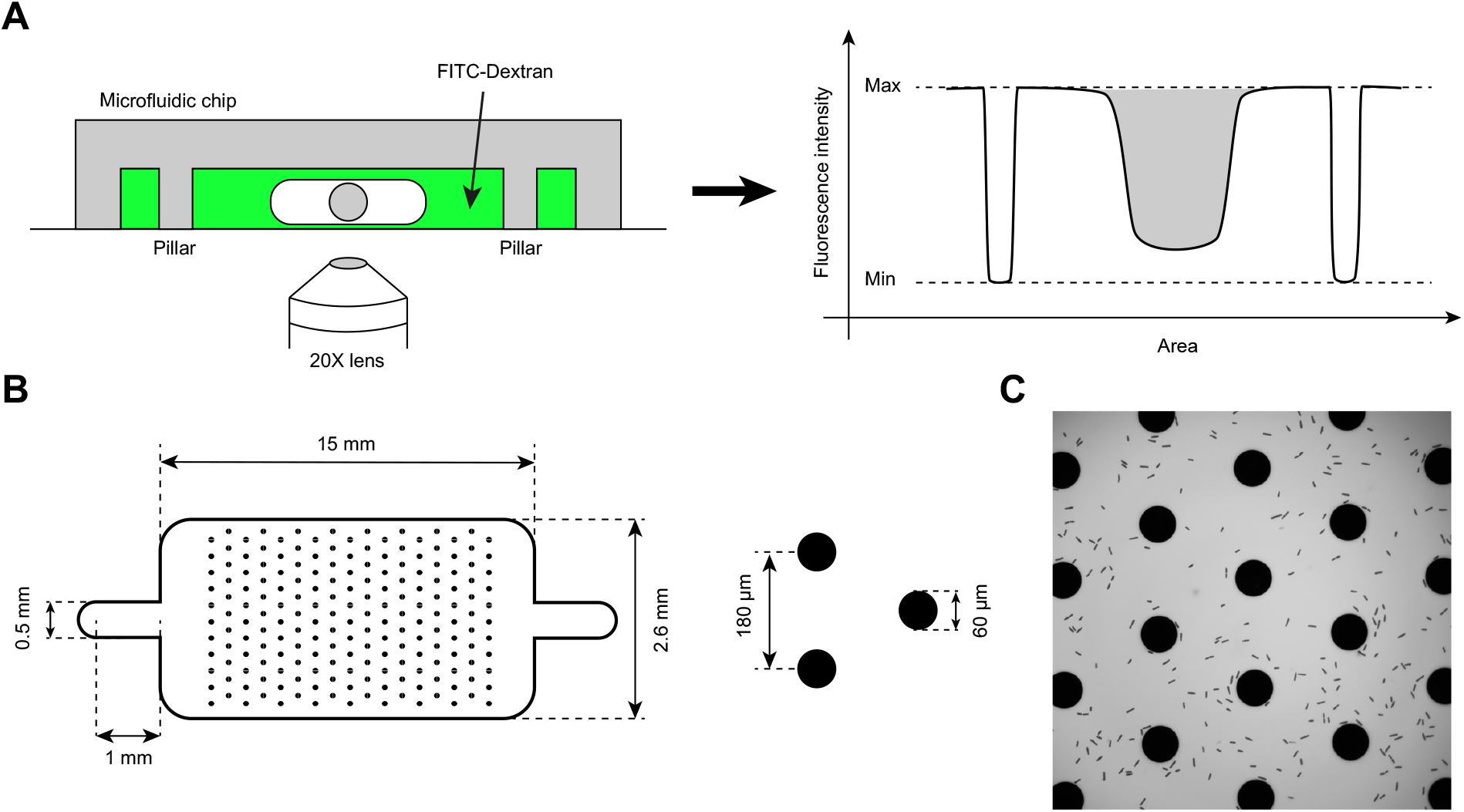
Microfluidic chips for FXm. **A.** Schematic of the principles of FXm (Cadart et al., 2017). Left: cells are injected in a FXm chip (see *B*) in the presence of FITC-Dextran. Pillars are integrated for ceiling support and image normalization. A low-magnification objective is used to acquire single-plane images (see *C*). Right: After normalization, the volume of the cell can be determined from the loss of fluorescence (grey area, the precise internal height of the chip must be determined (Cadart et al., 2017)). **B.** Schematic of the chamber used for fission yeast. Left: design and dimensions (schematic not to scale). Right: pillars are 60 μm in diameter and separated by 180 μm. **C.** Image of fission yeast cells in an FXm experiment (prior to image normalization).

Here we use the fission yeast *Schizosaccharomyces pombe* as a model system and demonstrate that the fluorescence exclusion method is a unique tool for measuring the volume of small cells. We engineer specific microdevices for yeast cells, assess the reproducibility and sensitivity of the system, and provide an improved way to perform large-scale analyses. Furthermore, we demonstrate the compatibility of the approach with multi-color imaging for evaluating single-cell volume in specific subpopulations. We then show how time-lapse experiments using these devices is a powerful strategy to couple yeast genetics with volume measurements for deciphering growth dynamics and size modulation. Interestingly, our results suggest that the use of FXm to re-evaluate previous models will be essential for understanding how cell volume is regulated in fission yeast, with implications for the general principles that govern cell size control in complex eukaryotes.

## RESULTS

### Microfluidic devices for determining yeast cell volume by FXm

FXm was developed to measure the volume of mammalian cells despite their complex morphologies (Cadart et al., 2017; Cadart et al., 2018). This method is simple and potentially compatible with any cell type, making it an ideal approach for investigating the regulation of cell volume in genetically amenable models such as yeast. However, to ensure an optimal dynamic range for accurate volume measurements, the height of the microfluidic chamber must be of the same order of magnitude as that of the cells. Indeed, the exclusion of fluorescent molecules in a microsystem several times higher than the cell would only marginally impact the total fluorescence intensity in the region of interest, affecting the reliability of the results. Microdevices employed to determine mammalian cell volume are therefore not adapted to small fission yeast cells, which require chambers of ~4-7 μm in height, depending on the strain studied. Interestingly, evaluation of FXm suggests that such reduced heights represent an advantage for the accuracy of these measurements (Model, 2020).

This design constraint led us to build specific and optimized devices for our yeast model. In particular, we determined a density of normalization pillars that is high enough to prevent the collapse of the chambers without significantly reducing the throughput of the method. Indeed, cells in close vicinity of these pillars are excluded from the FXm analyses due to the bias introduced by the presence of neighboring structures (Cadart et al., 2017). For this, we tested chambers with inter-pillar distances of 120, 180 and 270 μm (Fig. 1B). We found a distance of 180 μm between pillars coupled with an optimized cell concentration and preparation procedure to be the most reliable combination for the measurement of high numbers of individual cells in a single frame while limiting cell aggregation (Fig. 1C, see Materials and Methods).

### Measuring the volume of individual fission yeast cells using FXm

The necessity of using microchips of reduced height may represent a challenge for precisely measuring the volume of individual yeast cells. Indeed, in these conditions, the difference between the minimum (pillar) and maximum (no cell) fluorescence intensities (Fig. 1A), as well as the extent of fluorescent molecule displacement in the presence of a cell, are inherently low. This may render the technique less sensitive and more susceptible to noise, with consequences for its accuracy and reproducibility.

To evaluate whether FXm provides high quality results for yeast, we measured the size of wild-type fission yeast cells, comparing both the median volumes and volume distributions from independent experiments (see Materials and Methods). To this end, we used a 5.53 μm high chamber design and analyzed five replicates performed in separate devices built from the same master mold. Remarkably, we obtained highly reproducible data, with a median volume for the pooled datasets of 104.8 ± 24.5 μm^3^ (standard error for the 5 replicates, s.e.m.: 1.7 μm^3^), and comparable population profiles (Fig. 2A). This value for wild-type cells is consistent with previous results established by complex image processing (Baybay et al., 2020), validating FXm for cell size monitoring in yeast. In contrast to size at division, which only focuses on a small subpopulation, our method describes the volumes of the cells over the entire population, from birth to division. It therefore integrates the changes in size that occur during the division cycle. In addition, our experiments suggest that several repeats may be needed when evaluating median volume differences of less than 10 %, as such alterations are in the range of the observed experimental variability (Fig. 2A). When considering a geometric cell model consisting of a cylinder with half spheres at both ends and a constant diameter of 4 μm, an increase of 10 % in volume at division is the result of an increase in cell length at division from 14 to ~15 μm. Conversely, a similar change in cell volume at constant length is brought about by an increase in diameter of ~0.2 μm. In both cases, these differences are at the limit or below the resolution of the strategies routinely used for measuring cell length or width in *S. pombe*. This demonstrates the high sensitivity of FXm for yeast cell volume measurement, even when applying a conservative threshold of 10 % change.

**Figure 2.**
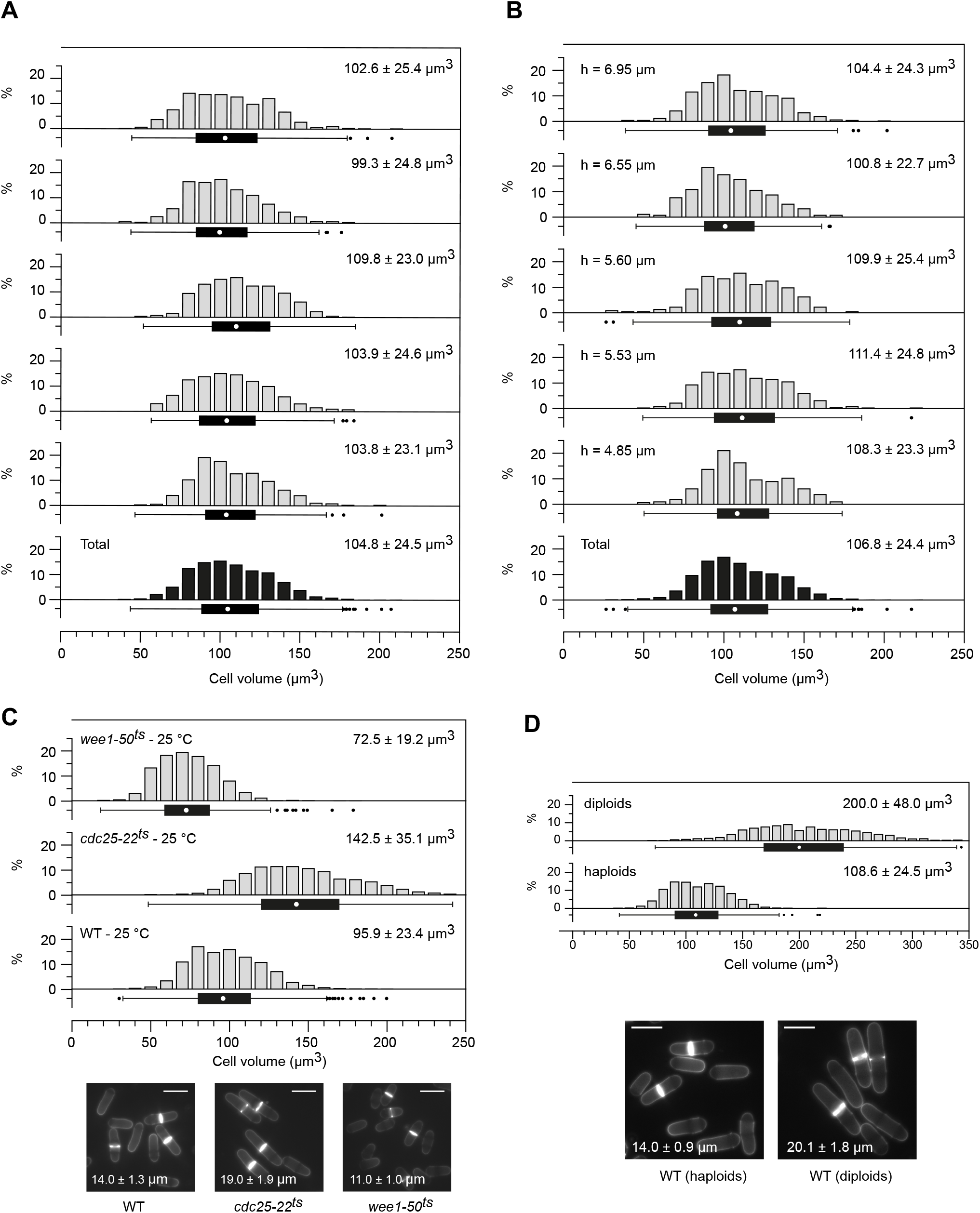
Measuring fission yeast cell volume by FXm. *A-D*, graphs are histograms with box and whiskers plots, indicating the min, Q1, Q3 and max, with outliers determined by 1.5 IQR (interquartile range). White dot indicates the median volume. Median volumes with s.d. are shown. **A.** Measurements of wild-type fission yeast in five independent experiments (grey) and the total pooled dataset (black). Chamber height: 5.53 μm; *n*≥316 for each replicate, *n*=2266 for the total dataset; values outside of the plot range (*n*\*) were excluded: *n*\*≤1 for each replicate. **B.** Measurements of wild-type cells from a single culture using five distinct devices of the indicated heights (grey) and the total pooled dataset (black). *n*≥225 for each replicate, *n*=1777 for the total dataset; *n*\*=0 for each replicate. **C.** Top: wild type (WT), *wee1-50^ts^* and *cdc25-22^ts^* temperature-sensitive strains were grown at the permissive temperature of 25 °C and their volumes measured by FXm. Pooled datasets (*n*≥1144) of three independent experiments (*n*≥337 for each replicate, Fig. S1A) are shown. *n*\*≤6 for each replicate. Bottom: blankophor images with cell length at division (averages with s.d. for pooled datasets of three independent experiments, *n*≥100 for each replicate). Scale bars = 10 μm. At 25 °C, WT cells have a reduced volume and cell length at division compared to 32 °C (see *A*, *B* and Figs. 4-7). **D.** Top: comparison of volume distributions in populations of WT haploid and diploid fission yeast cells by FXm. Cells were grown in supplemented EMM6S at 32 °C. Pooled datasets (*n*≥1073) for three independent experiments (*n*≥207 for each replicate, Fig. S1B) are shown. *n*\*≤1 for each replicate. Bottom: blankophor images with cell length at division of the corresponding strains (averages with s.d. for pooled datasets of three independent experiments, *n*≥100 for each replicate). Scale bars = 10 μm. In EMM6S, WT cells have a volume that is similar to that in EMM (compare with *A*, *B* and Figs. 4-7) but a reduced cell length at division (compare with Fig. 4A, 5A and 7), suggesting a difference in cell width between these conditions.

Next, as molds deteriorate when heavily used and given that standard microfabrication procedures make it difficult to generate highly reproducible chamber heights, we set out to evaluate how changes in the dimensions of the devices may impact volume determination. We therefore produced microchips of various heights and measured the size of wild-type fission yeast cells by FXm. Again, we observed remarkably similar median volumes and volume distributions between measurements (Fig. 2B – median volume of the pooled datasets: 106.8 ± 24.4 μm^3^; s.e.m. for the 5 replicates: 1.9 μm^3^). This shows that data obtained with different devices can be reliably compared.

Finally, we further validated the approach by assessing the volumes of known fission yeast size mutants as well as wild-type diploid cells. In particular, we measured the size of well-described temperature-sensitive cell cycle mutants: *wee1-50^ts^* (small cells) and *cdc25-22^ts^* (large cells) (Fantes, 1979; Fantes and Nurse, 1978; Nurse, 1975). As anticipated, *wee1-50^ts^* and *cdc25-22^ts^* cells grown at the permissive temperature of 25 °C showed significantly smaller and larger volumes than wild type, respectively (Fig. 2C, Fig. S1A). Note that the apparent broader distribution in *cdc25-22^ts^* results from increased overall size, as the volume profiles of these strains are similar when the data are normalized to their respective median sizes (Fig. S1C). In the case of wild-type diploid cells, we found that the median population volume is 1.84-fold higher than for haploid cells (Fig. 2D, Fig. S1B).

Altogether, these experiments demonstrate that FXm with specifically adapted microfluidic chips is compatible with the analysis of yeast cell volume and produces highly reproducible results. Furthermore, in contrast to measurements of cell length and width, FXm does not rely on 2D parameters that are difficult to accurately determine. In particular, it will allow for identifying changes in volume that result from limited alterations in cell diameter. Finally, our data also set a first conservative threshold of 10 % for the estimation of differences in cell volume between populations.

### Improving FXm image analysis

FXm image analysis consists of three steps (Cadart et al., 2017): image normalization, cell selection and volume calculation. However, as FXm requires a low magnification objective, the size of yeast cells and the high number of cells in a frame make it tedious to manually select individual cells using the code established for mammalian cells. Interestingly, the normalization procedure generates a mask that separates pillars and cells from the background (Cadart et al., 2017). Since the majority of objects extracted at this step corresponds to valid individual cells, we took advantage of this mask and developed a Python script to assist the user in cell selection (see Supplementary information). Thus, instead of manually delineating the multiple regions of interest (Fig. 3A), this tool offers two distinct strategies.

**Figure 3.**
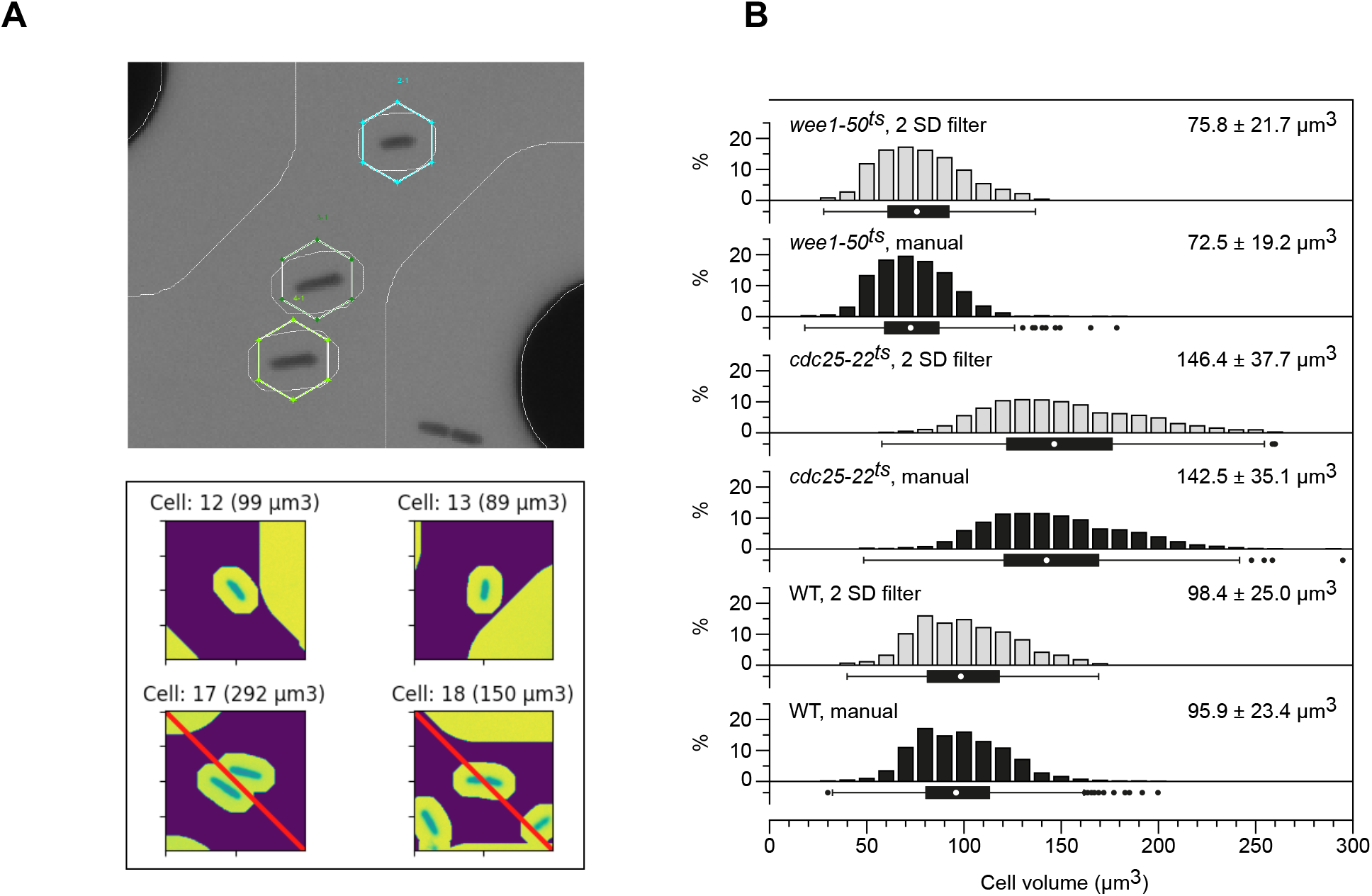
Analysis pipeline for FXm data. **A.** Top: representative image showing the selection of individual cells using the original FXm analysis code. Each cell must be identified from the FITC image (see Fig. 1C; the area shown has been magnified for display purposes) and manually selected. Bottom: representative image showing the tool that we developed, taking advantage of the normalization mask for cell selection. All objects are automatically identified and displayed at a higher magnification. Cell aggregates (bottom left), recently-divided cells (bottom right) or artifacts can be excluded by the user (red-barred). **B.** Comparison of FXm results using manual (black) *vs*. automated (grey, 2 s.d. filter) cell selection. Data for manual selection are from Fig. 2C. For automated selection, 1) the same images as for the manual method were used, pooling the data from all replicates, and 2) a filter was applied that excludes entries that are 2 s.d. above and below Q3 and Q1 (*n*\*\*), respectively (unfiltered selected objects: *n*≥1390, excluded objects *n*\*\*≤200). The overlap between the two approaches for both correctly assigned cells and outliers is between 87 and 92 %. Graphs are as in Fig. 2.

First, a manual cell selection mode can be chosen. Cells pre-selected by the normalization mask are magnified and displayed (Fig. 3A), allowing the user to rapidly exclude incorrect objects such as dust particles or cell aggregates. This approach makes the analysis of a high number of cells easier and faster. Second, an automated and non-discriminatory selection can be applied. To exclude aberrant measurements (*e.g*. cell aggregates, small dead cells), thresholds are then automatically set to remove outliers based on the standard deviation (s.d.) of the population data. This is more appropriate than the use of absolute cut-offs, which may need to be adjusted from strain to strain. After testing different s.d. thresholds (data not shown), we found that excluding values outside of 2 s.d. (1 IQR for a normal distribution) for the first and third quartiles was optimal for obtaining results similar to those established with the manual method, correctly selecting between 87 and 92 % of the objects in our experiments (Fig. 3B). Finally, our script also provides the option of separating the selected cells in different groups. This is particularly useful when determining the volumes of distinct subpopulations, for instance based on additional fluorescence markers (see below).

Collectively, the improvements brought by our analysis code facilitate FXm and provide more options to the user for complex studies of cell populations. Interestingly, while manual selection is the most accurate approach, we show that automated analysis with outlier exclusion allows a rapid assessment of high numbers of cells while remaining sufficiently reliable for screening the volumes of many different strains.

### Small variations in cell dimensions and population heterogeneity

Our initial volume measurements in wild-type fission yeast cells and the evaluation of FXm experimental variability suggest that changes in volume ≥10 % can be reliably detected (Fig. 2). To further assess the sensitivity of FXm, we first determined the volume of cells showing a marginal difference in length at division compared to wild type. To this end, we took advantage of a fission yeast strain whose proliferation relies on a minimal cell cycle control network (MCN). In this background, cell proliferation is solely dependent on the oscillation of a single cyclin-dependent kinase (CDK) activity between two thresholds (high for mitosis and low for S phase) (Coudreuse and Nurse, 2010). This is achieved through the expression of a fusion protein between cyclin B/Cdc13 and Cdk1/Cdc2, in the absence of all other cell cycle cyclins (Coudreuse and Nurse, 2010). Interestingly, while behaving similarly to wild type (Coudreuse and Nurse, 2010), MCN cells show a minor increase of ~7 % in their length at division in our experimental conditions (15 μm for wild type and 16.1 μm for MCN cells, Fig. 4A). This is predicted to contribute to a similar increase in volume according to the geometric model of *S. pombe* cells (see above). Unexpectedly, FXm measurements showed a median volume difference of 17 %between these strains (Fig. 4A, Fig. S2; compare WT and MCN). This suggests that the change in size between wild type and MCN cells is not solely the result of an increase in cell length. Importantly, it demonstrates that FXm is a promising approach for investigating volume regulation in yeast, providing novel and more in-depth information on cell size and geometry.

**Figure 4.**
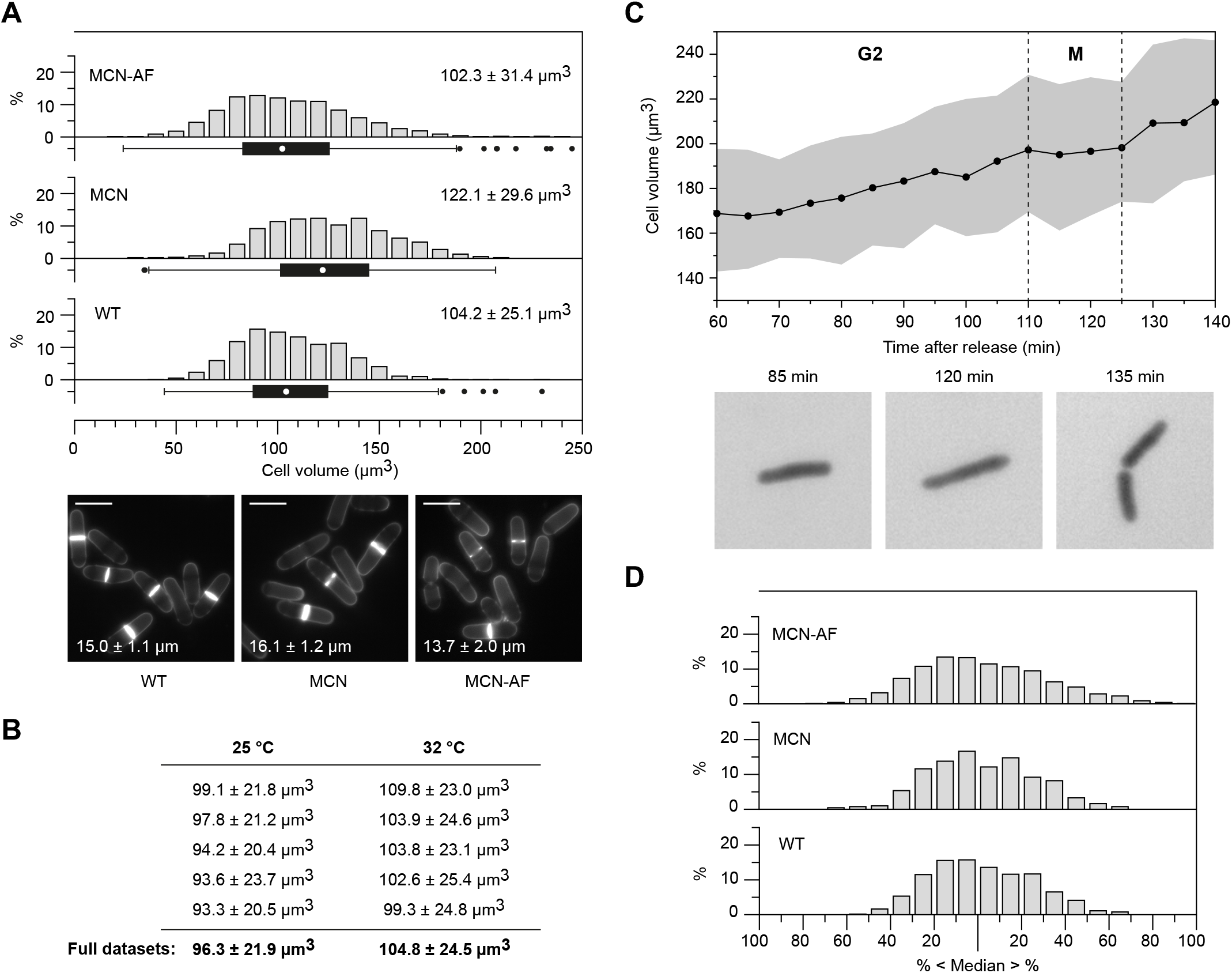
FXm allows for the detection of subtle changes in cell volume and size distribution. **A.** Top: volume measurement of WT, MCN and MCN-AF cells by FXm. Pooled datasets (*n*≥913) of three independent experiments (*n*≥172 for each replicate – Fig. S2) are shown. *n*\*≤1 for each replicate. Graphs are as in Fig. 2. Bottom: blankophor images with cell length at division of the corresponding strains (averages with s.d. for pooled datasets of three independent experiments, *n*≥100 for each replicate). Scale bars = 10 μm. **B.** Comparison of the median volumes for 5 independent replicates of WT at 25 and 32 °C. Full datasets are in Fig. S3A. Difference in median cell volume: 8.8 %. **C**. MCN cells grown in EMM6S at 32 °C were G2-arrested using 1 μM 3-MBPP1 for 160 min and allowed to synchronously re-enter the cell cycle by washing off the inhibitor (Coudreuse and Nurse, 2010) (Fig. S3B). Cells were then injected in a FXm chip, and images were acquired at 5 min intervals throughout the next G2, after cells had undergone a first round of mitosis and cell division (60 min, Fig. S3C). Top: median population volumes at the indicated time points (black line – automated cell selection) with IQR (grey area) are shown. Note that the higher volume for MCN cells compared to other Figures results from the initial G2 block, during which cells grow without dividing. The phases of the cell cycle during which cell volume was measured (first G2 and mitosis/M after release – Fig. S3C) are indicated. G2 growth rate: 0.59 μm^3^.min^-1^ (linear regression from 60 to 110 min, prior to the mitotic plateau). From 125 to 140 min, the automated selection does not separate newly divided cells (17.4%, 25.9% and 25.5% of the selected objects at 130, 135 and 140 min, respectively, are pairs of cells; the remainder of the cells have not undergone cytokinesis at these time points). This results in an apparent increased growth rate. Note the delay in cell cycle progression compared to Fig. S3C, which may be due to the growth conditions in the absence of medium flow (see Fig. 8B). For each time point, 214≤*n*≤278 (unfiltered automatically selected objects). Outliers were removed using the 2 s.d. filter (see Fig. 3B; 28≤*n*\*\*≤49, representing a maximum of 18% of the number of selected objects). Bottom: representative images for cells in FXm chips at 85 min (G2), 120 min (M) and 135 min (pairs of sister cells represent a significant subset of the population at this time point, see above). **D.** Volume distribution for the indicated strains as a percentage of the median volume of the population. Data are as in *A*. *n*\*≤10.

Next, we tested whether FXm can reliably reveal volume changes below our initial threshold of 10 %. First, building on the results in Fig. 2A and 2C, we showed that a difference in volume of less than 10 % between wild-type cells grown at 32 *vs*. 25 °C could be robustly established by simply increasing the number of experimental replicates (Fig. 4B and Fig. S3A). We then investigated whether even smaller alterations in volume at the population level could be detected as cells progressively change size over time. As a proof-of-concept, we used MCN cells, as they harbor a mutation in the Cdc2 moiety of the fusion module that makes them sensitive to dose-dependent and reversible inhibition of CDK activity by non-hydrolysable ATP analogues such as 3-MBPP1 (Bishop et al., 2000; Coudreuse and Nurse, 2010). This allowed us to block cells in G2 using the inhibitor, let them synchronously re-enter the cell cycle upon inhibitor wash-off (Fig. S3B, C) and monitor volume changes throughout the following G2 with high time resolution (Fig. 4C). Remarkably, even considering the experimental variability observed at certain time points, very small increases in volume (<5%) and changes in growth rates could be measured, demonstrating the high sensitivity of FXm in this context.

Another critical parameter in our understanding of cell size homeostasis is the size distribution in the population. This reflects the cell-to-cell heterogeneity in cell cycle progression and the strength of the size control operating in these cells. To date, most studies in fission yeast have focused on variability in cell length at division. However, not only does this approach exclude the majority of cells, but our results above suggest that evaluating cell volume distribution may lead to different conclusions. To establish whether FXm is sufficiently sensitive for monitoring alteration of cell size profiles, we took advantage of MCN cells in which the target residues of the conserved Wee1/Cdc25 feedback loop on Cdc2 are mutated (T14A Y15F, MCN-AF) (Coudreuse and Nurse, 2010). Loss of this mitotic switch results in an increase in cell-to-cell variability in length at division (Coudreuse and Nurse, 2010). In the non-supplemented medium used in our experiments, we also found that these cells are on average slightly smaller than wild type (Fig. 4A). As anticipated, we observed a broader distribution of cell volume in MCN-AF compared to MCN and wild-type cells (Fig. 4A, D). Interestingly, the relative changes in median volume *vs*. average length at division between MCN-AF, MCN and wild type provide insights into the dimensions that are altered in these strains. (Fig. 4A, Table S1): while MCN-AF and MCN appear to differ only in length (same ratios for volume *vs*. size at division), both strains are likely to show an increase in cell diameter compared to wild type (differing ratios).

Altogether, these results demonstrate that FXm is easier and more sensitive than previously used methods, enabling the monitoring of small alterations in cell volume and changes in size homeostasis in populations of yeast cells. Coupling these data with more traditional cell length measurements will shed light on how changes in volume are mediated and whether modulations of cellular geometry occur.

### Measuring the size of cells with altered geometries and morphologies

FXm allows for measuring small changes in size that are difficult to accurately evaluate from cell length and diameter measurements. The use of length as a proxy for fission yeast size is even more problematic when studying mutants or conditions where dimension scaling or cell shape is altered. Furthermore, strategies using cell outline detection to extrapolate cell volume may not be sufficiently reliable, given the influence of minor changes in diameter on cell volume.

In this context, we determined the volumes of various strains showing defects in morphology and shape. First, we assessed the size of strains lacking Rga2, Rga4 or Rga6, which are members of the RhoGAP family. These factors are involved in polarized growth, and their loss impacts cell geometry (Das et al., 2007; Revilla-Guarinos et al., 2016; Soto et al., 2010; Villar-Tajadura et al., 2008). When ranking these strains according to their length at division, we found that Δ*rga2* cells are the longest, followed by Δ*rga6* and Δ*rga4* (Fig. 5A). Strikingly, FXm, which is unaffected by changes in either morphology or geometry, led to the opposite conclusion, with Δ*rga4* and Δ*rga2* cells having the largest and smallest volumes, respectively (Fig. 5B, Fig. S4). This shows that cell length measurement is not reliable as a proxy for cell size when studying such mutants. Our data also demonstrate the advantages of FXm for establishing size differences between strains and determining the influence of diverse pathways on cell volume regulation.

**Figure 5.**
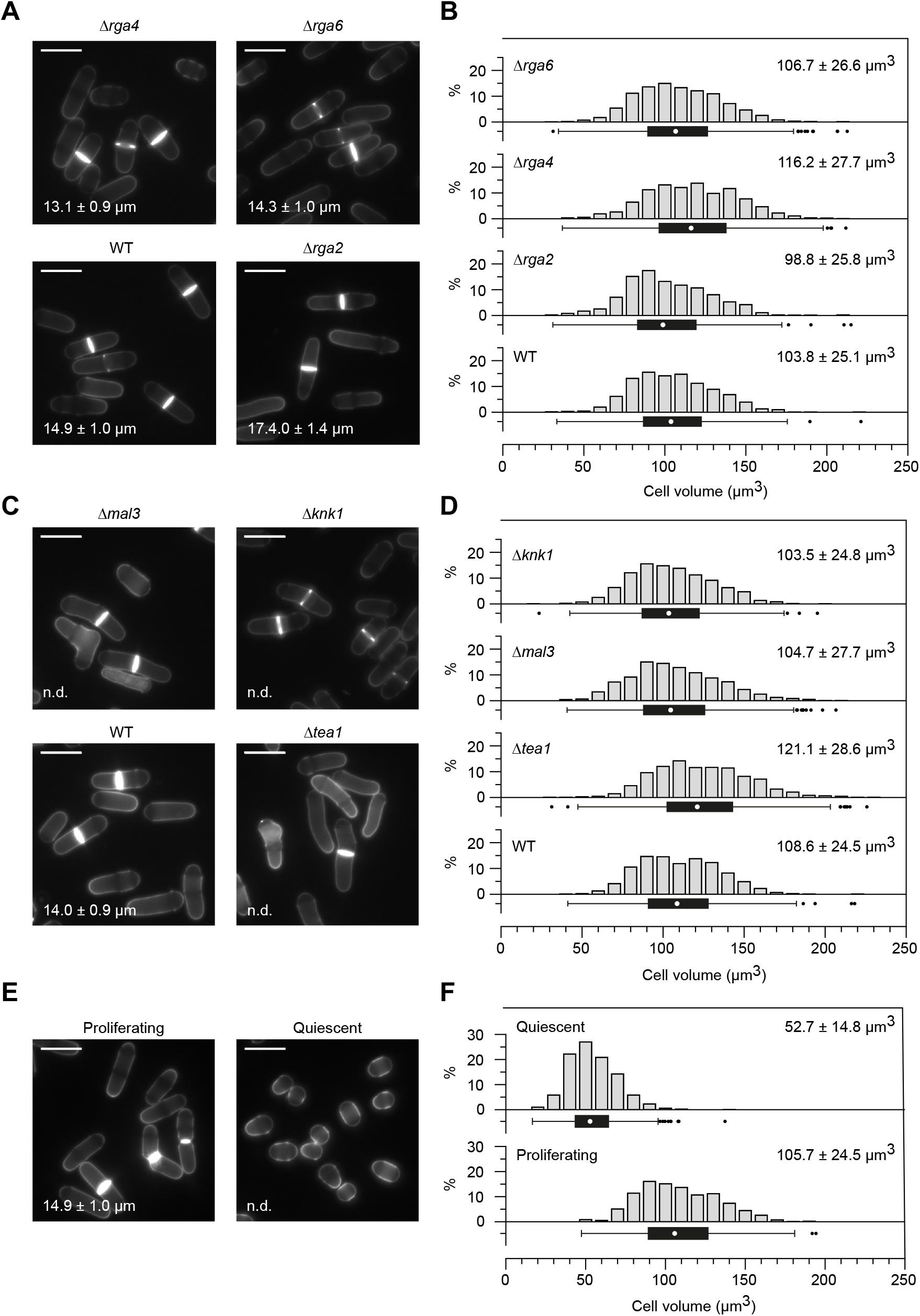
Measuring the volume of yeast cells with altered morphologies. **A, C, E.** Blankophor images with cell length at division of the indicated strains (averages with s.d. for pooled datasets of three independent experiments, *n*≥100 for each replicate). n.d.: not determined. Scale bars = 10 μm. **B.** Volume measurement by FXm of the strains in *A*. Pooled datasets (*n*≥832) of three independent experiments (*n*≥268 for each replicate – Fig. S4) are shown. *n*\*≤1 for each replicate. **D.** Volume measurement by FXm of the strains in *C*. Pooled datasets (*n*≥940) of three independent experiments (*n*≥230 for each replicate – Fig. S5) are shown. *n*\*=0 for each replicate. Data for WT are as in Fig. 2D (haploids). **E.** Blankophor images of WT cells in exponential growth and quiescence. For quiescent cells, samples were taken 24 h after the culture reached OD_595_ 0.4. For proliferating cells, cell length at division is shown (average with s.d. for pooled dataset of three independent experiments, *n*≥100 for each replicate). n.d.: not determined. Scale bars = 10 μm. **F.** Volume measurement by FXm of the cells in *E.* Pooled datasets (*n*≥965) of three independent experiments (*n*≥238 for each replicate – Fig. S6) are shown. *n*\*=0 for each replicate. *C*, *D*: Experiments were carried out in EMM6S due to the presence of auxotrophies in some of the strains (Table S2). *B*, *D*, *F:* Graphs are as in Fig. 2.

Next, we used FXm to evaluate the size of cells for which length can be difficult to measure or inherently inaccurate due to morphological alterations. To this end, we used bent cells (Δ*knk1*, Δ*tea1*) as well as cells with significant shape defects and irregular diameters (Δ*mal3*) (Beinhauer et al., 1997; Mata and Nurse, 1997; Scheffler et al., 2014) (Fig. 5C). As anticipated, while cell length at division could not be determined without excluding a significant fraction of the cells, FXm allowed us to establish the profiles of cell volume in these populations and compare them with that of wild-type cells. This again highlights the power of this method for studying size control mechanisms independently of cell morphology (Fig. 5D, Fig. S5). Interestingly, in cells with morphological defects, both the median volume and size distribution may change (Figs. S4, S5), potentially reflecting alterations of their cell cycle organization and associated growth patterns. This opens the door to investigating how cell shape may delineate the complex interplay between cell growth and the division cycle at the single-cell level.

Finally, we evaluated FXm data for significantly smaller cells that have stopped dividing. Upon glucose exhaustion, fission yeast cells exit the cell cycle and enter a non-proliferative quiescent state, which is associated with a dramatic reduction in cell size. Given the shape of the cells in these conditions (Fig. 5E), determining their length and width to extrapolate their volume is difficult and unreliable. In contrast, FXm established high-quality profiles of cell volume in these populations (Fig. 5F, Fig. S6). This will allow for investigating the impact of cell size on cell physiology and aging in this critical cellular state.

Collectively, this set of experiments demonstrates the versatility of FXm, which provides an unprecedented ability to measure the size of any strain, irrespective of cell morphology and physiological status. Our results also suggest that previous conclusions on size control in fission yeast may need to be re-evaluated using FXm.

### Measuring the volume of subpopulations

All experiments presented so far focus on the determination of cell volume in entire cell populations. However, extracting the size of cells in specific subpopulations is critical for understanding how different biological processes may interact with the mechanisms underlying cell size control and how cell growth and proliferation are coupled. Thus, associating FXm with the use of other intracellular fluorescent markers would represent a powerful method to take advantage of yeast genetics and explore the complexity of size regulation in different conditions and throughout the cell cycle. Similarly, this could allow for discriminating between distinct strains expressing specific markers in a complex population, opening the door to exploring non-cell autonomous processes that may feed into volume regulation.

To evaluate these approaches, we set out to determine cell volume at mitosis, using the non-histone chromatin-associated protein Nhp6 coupled to the red fluorescent marker mCherry. First, we validated that the presence of Nhp6::mCherry in the nucleus did not induce any significant signal in the FITC channel under FXm conditions (Fig. S7A). This also showed that the slightly reduced volume of these cells compared to wild type (Fig. 6A, Fig. S7B) results from an alteration of Nhp6 function due to the tag rather than an experimental bias from mCherry fluorescence. Using our improved FXm code, we segmented the population of binucleated cells (~11 % of the total number of cells) and determined their median volume (126.3 ± 18.2 μm^3^, compared to 93.5 ± 21.3 μm^3^ for mononucleated cells, Fig. 6B). The partial overlap between mono- and binucleated cells is consistent with the interruption of growth during mitosis: cell size does not change throughout nuclear division, and growth only resumes in G1. Note that the volume of binucleated cells measured in our assay is consistent with previous calculations of cell volume at division (Nurse, 1975) but significantly differs from those presented in a more recent study (Navarro and Nurse, 2012). This further demonstrates that fission yeast cell size cannot be reliably and reproducibly determined using cell length and diameter. Altogether, our experiments validate the possibility of combining FXm with additional markers in yeast for more complex investigations of cell size control and dynamics.

**Figure 6.**
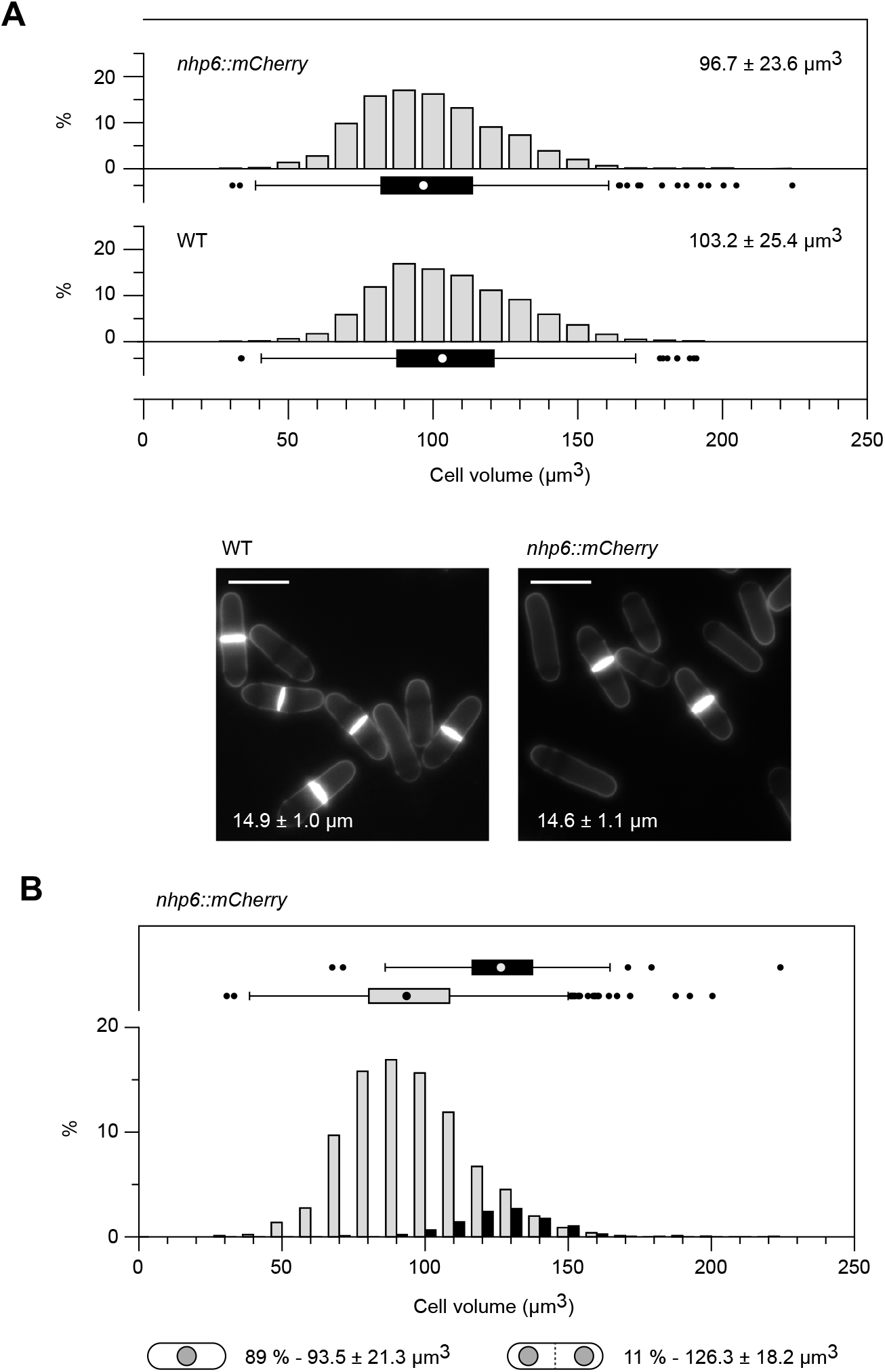
FXm is compatible with the use of fluorescent markers to study subpopulations. **A.** Top: volume measurement of WT and cells expressing *nhp6::mCherry* by FXm. Pooled datasets (*n*≥1568) of three independent experiments (*n*≥499 for each replicate – Fig. S7B) are shown. *n*\*=0 for each replicate. Median volume values with s.d. are shown. Bottom: blankophor images with cell length at division of the strains above (averages with s.d. for pooled datasets of three independent experiments, *n*≥110 for each replicate). Scale bars = 10 μm. **B.** Volume comparison between mono- (grey) and bi-nucleated (black) cells in the population of *nhp6::mCherry* cells as in *A*. The percentages and median volumes for the mono- (*n*=1619) and binucleated (*n*=196) subpopulations are indicated. Binucleated cells include both septated (dashed line) and non-septated cells. *A*, *B*: Graphs are as in Fig. 2.

### Dynamic changes in fission yeast cell volume at the population and single-cell levels

Next, we tested the use of FXm for assessing dynamic changes in yeast cell size. First, we ascertained whether coupling time course experiments with FXm allows for following induced alterations in cell volume over time using asynchronous populations. A wide range of growth conditions (*e.g*. heat stress (Vjestica et al., 2013), osmotic stress (Millar et al., 1995), change in nutrient availability (Fantes and Nurse, 1977; Petersen, 2009)) as well as experimental modulation of specific cellular pathways can trigger cell size changes. Understanding the dynamics of such changes may be key to determining the implication of these processes in cell size homeostasis. To evaluate FXm in this context, we took advantage of the analog-sensitivity of MCN cells. Indeed, reducing Cdc2/CDK activity in fission yeast using this method leads to an increase in cell size (Chen et al., 2016; Coudreuse and Nurse, 2010). We therefore measured cell volume in asynchronous MCN cells at 40-minute intervals (~1/4 of the cell cycle) after treatment with 0.05 μM 3-MBPP1 (Fig. 7). Our data show that MCN cells progressively increase their volume, up to 1.7-fold after one doubling time. Strikingly, comparison of the volume *vs*. size at division ratios throughout the experiment (Table S1) suggests complex dynamics in the respective modulations of length and diameter upon CDK inhibition. This demonstrates that FXm 1) can be used to follow short-term alterations in median cell size and size distribution in asynchronous yeast populations and 2) provides new perspectives for our understanding of how the geometrical dimensions of these cells contribute to cell volume.

**Figure 7.**
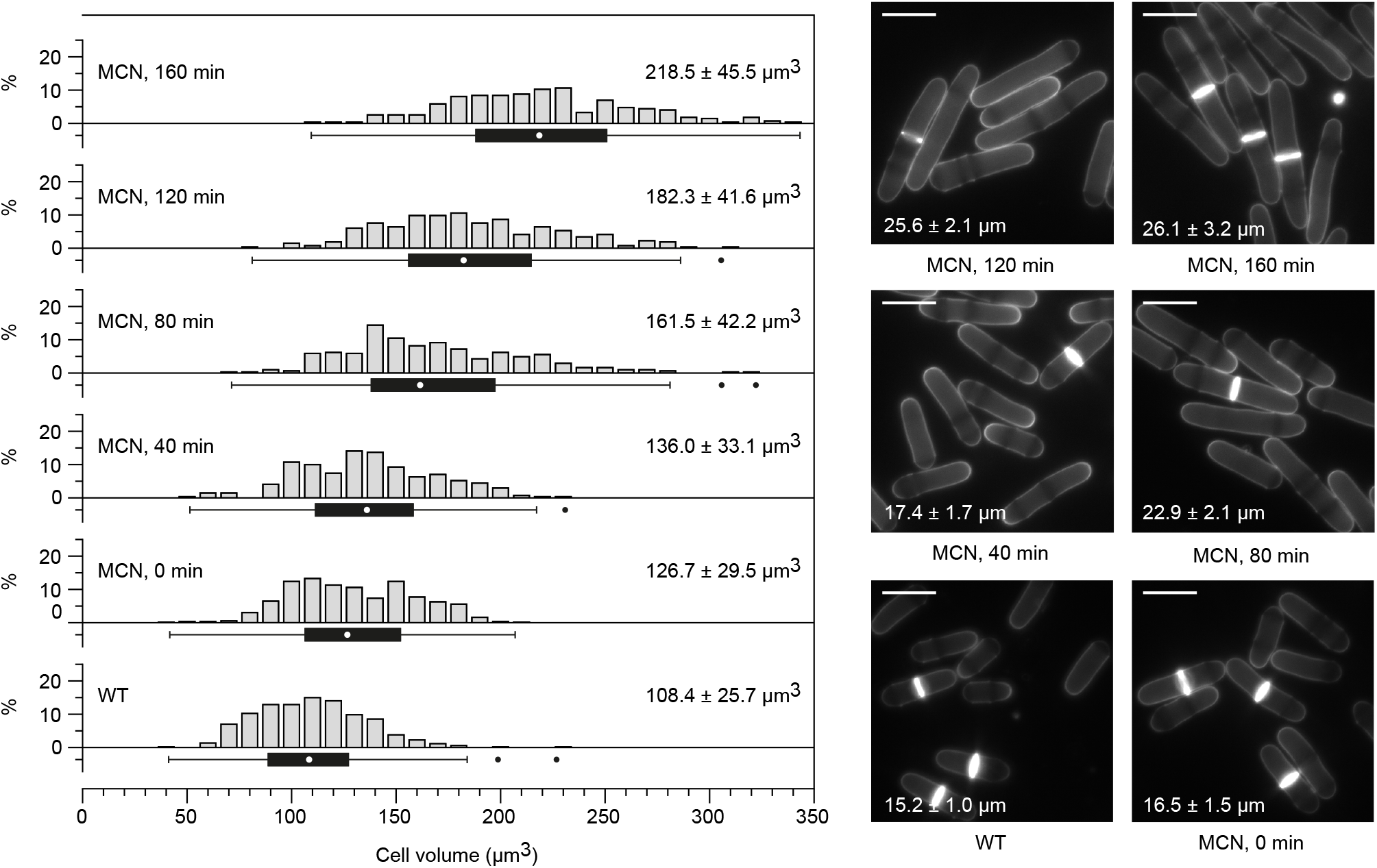
Dynamic changes in cell volume at the population level. Left: FXm volume measurement of MCN cells after treatment with 0.05 μM 3-MBPP1. Untreated WT cells were used as a control. For MCN, samples were collected every 40 min after addition of the inhibitor (*n*≥265 at each time point, *n*=528 for WT). *n*\*≤2 for each dataset. Graphs are as in Fig. 2. Right: blankophor images with average cell length at division and s.d. for the experiment in *A*. *n*≥100 for WT and MCN (0, 120 and 160 min). For the 40 and 80 min time points, the initial delay in mitosis due to the inhibitor treatment leads to a reduction in the number of dividing cells, making it difficult to obtain similar numbers for the calculation of cell length at division (*n*=45 and 22, respectively). Septation indices: WT: 11.5 %, MCN 0 min: 20.4 %, MCN 40 min: 1 %, MCN 80 min: 2 %, MCN 120 min: 15.5 %, MCN 160 min: 26.9 %; *n*≥400 for each time point. Scale bars = 10 μm.

We then asked if FXm could be applied to analyze individual yeast cells in time-lapse experiments. First, we determined the experimental noise for single cells in our chips by comparing 20 successive measurements of 10 individual cells over a period of ~6 seconds (Fig. 8A). Making the assumption that no volume changes occur in such a short time, this showed that the variability inherent to FXm when using chips of reduced height remains very limited. Interestingly, it also suggests that when monitoring single cells, volume alterations as low as ~3 % can be detected. Next, to perform time-lapse experiments of several hours, we began by using modified chips in which the cell injection inlets were made bigger (from 2.5 to 4 mm), acting as nutritional reservoirs to maintain growth conditions as stable as possible. As fission yeast cells are non-adherent, this also required pre-coating of the chip coverslips with lectin. In this context, we observed a linear increase in volume during each cell cycle and a plateau in size around the time of cell division, corresponding to the well-described growth arrest at mitosis (Mitchison, 1957; Mitchison and Nurse, 1985) (Fig. 8B). However, we found that in these experiments, cells divide at smaller sizes in succeeding cell cycles, suggesting that the built-in reservoirs are insufficient to ensure a favorable growth environment throughout our assay. We therefore applied a constant flow of fresh medium containing FITC-Dextran (~3-5 μL.min^-1^) for the entire duration of the experiments. In this setup, we did not detect any consistent reduction in cell volume at division after each cycle (Fig. 8C). Furthermore, in the presence of flow, the growth rate (see Materials and Methods) was significantly higher than without medium renewal (Fig. 8B, C).

**Figure 8.**
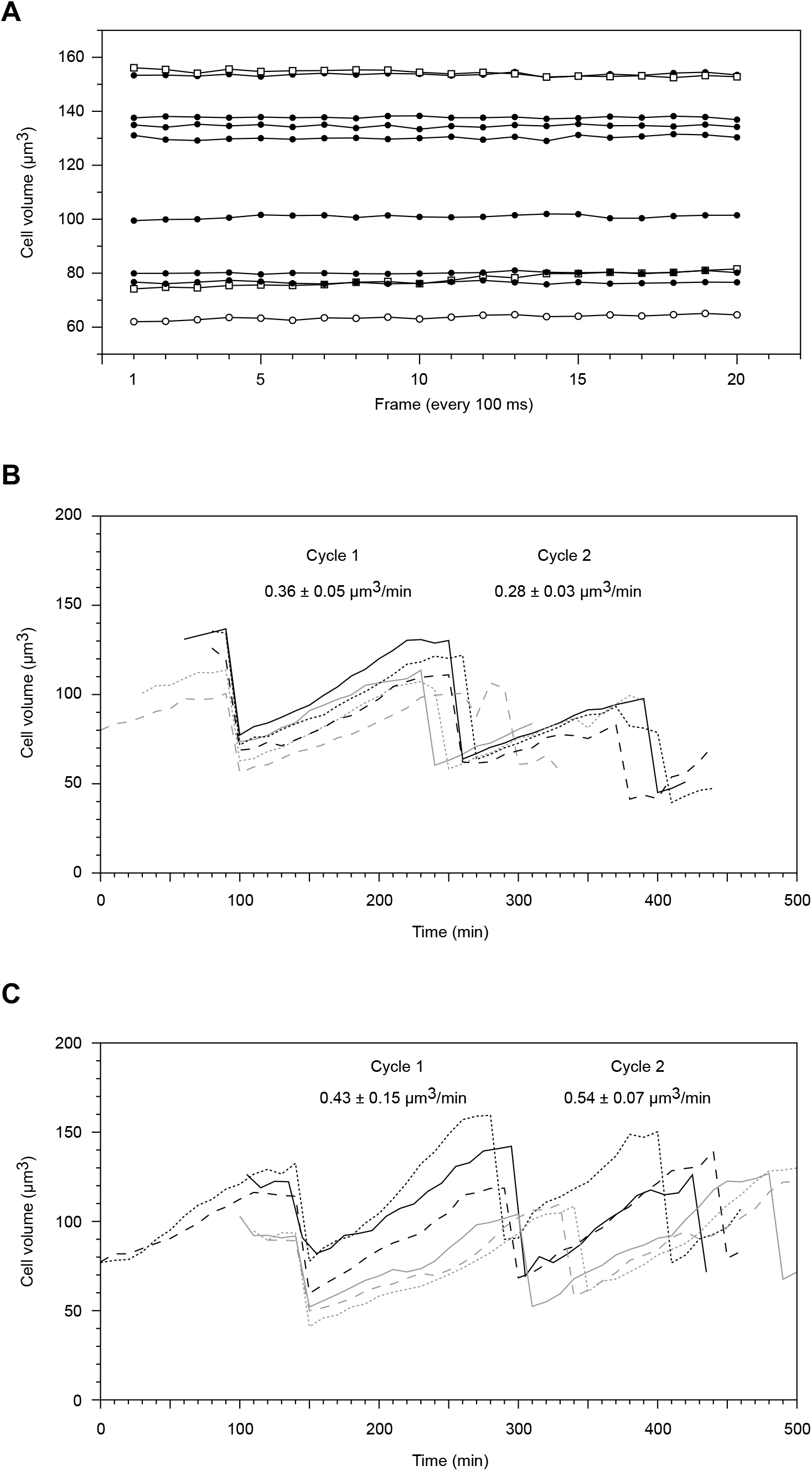
Determination and real-time monitoring of single-cell volume by FXm. **A.** 20 consecutive FXm images were acquired (see Materials and Methods; 100 ms exposure per image over a total time of 6 s) and the volumes of 10 individual cells were followed throughout the experiment. The generation time of WT cells in these conditions is ~140 min. The coefficients of variation are between 0.3 and 3.1%. **B.** FXm time-lapse experiment using WT in EMM at 32 °C without medium flow. **C.** FXm time-lapse experiment using WT in EMM at 32 °C with a constant flow of medium (3-5 μL.min^-1^) containing FITC-Dextran. *B, C*: individual traces represent the volume dynamics of single-cell lineages (at each division, only one of the two daughter cells is further monitored). Volumes were determined every 10 min. For display purposes, the traces are aligned to the first division (see Materials and Methods). Average growth rates with s.d. are shown for each cycle. Note that the growth rate in *C* is consistent to that obtained from the bulk analysis of a synchronized population (Fig. 4C).

These data demonstrate that FXm is a unique tool for monitoring volume dynamics of small cells with a high time resolution and for detecting changes in growth rate at the single-cell level. This will allow for investigating the immediate responses of cells to environmental perturbations and the behaviors of separate yeast cell lineages. In addition, this will make it possible to relate volume at birth, growth rate and volume at division for a given cell. Coupled with additional markers, this technique thus offers an unprecedented entry into the investigation of cell size control and its interplay with complex biological processes.

## DISCUSSION

Regulation of cell volume is critical for eukaryotic organisms, but the mechanisms underlying this process are still unclear. Yeast models have played a pioneering role in deciphering this complex trait and remain at the forefront of this very active research field. However, the difficulty of accurately measuring cell volume in these small unicellular organisms has been an obstacle for our understanding of size control and homeostasis. Here we show that the fluorescence exclusion method (FXm) is ideal for evaluating the volume of small yeast cells in diverse contexts. It provides high quality data on the volume profiles of yeast populations irrespective of cell morphology and physiological state, allows for single-cell analysis of volume changes with high time resolution and is compatible with the use of additional fluorescent markers. Our experiments also demonstrate that FXm gives researchers unique access to in-depth analyses of yeast cell volume dynamics and their coupling with cell cycle progression, integrating the size of cells at all stages of their division cycle. Furthermore, combining FXm with time-lapse imaging represents an unprecedented tool for exploring the changes in growth rate and cellular geometry that occur not only during proliferation or upon major transitions such as quiescence entry and exit, but also when cells respond to acute challenges. Thus, we believe that this technique will become a new standard for investigating cell size control in yeast, with implications for our knowledge of size regulation in complex eukaryotes.

Interestingly, our work led to a number of observations that may question existing models of cell size regulation. First, research on *S. pombe* size control and its interplay with cell cycle progression has mostly been based on the assumption that the diameter of proliferating haploid fission yeast cells is 1) constant from one cell to another and 2) homogenous along their growing axis. Our results clearly indicate that these assumptions can lead to incorrect conclusions when comparing strains, as shown using various mutants (Figs. 4, 5). In this context, FXm has the advantage of being a direct method for volume measurement that does not rely on any pre-conceived idea of cell geometry. Coupling FXm with cell length measurements also offers an additional level of understanding of cell size dynamics. For example, our data suggest that when cells are arrested in their division cycle through CDK inhibition, the geometric changes that occur over time are surprisingly complex, with length and volume alterations showing different kinetics (Table S1, MCN + 3-MBPP1). Similarly, we find that simplifying the architecture of cell cycle control or growing cells in distinct environments has unanticipated impacts on cell geometry, affecting both cell length and diameter (Table S1).

Moreover, focusing on subpopulations, as is the case when using cell length at division as a proxy for cell size, may hamper our comprehension of important processes linking cell growth, morphology and proliferation. For instance, the apparent changes in size distribution at the population level in morphology mutants suggest that the interplay between growth rate and cell cycle phases may be altered in these cells. This again may shed light on potentially novel inputs and mechanisms that modulate cell size dynamics. In this context, the different cell geometries of *rga* mutants (Table S1), as indicated by our comparisons of volume *vs*. cell length at division ratios, may represent a promising model for uncovering the key parameters used by the cells to “monitor” their size.

Our work demonstrates that previous models that describe the way cell size is regulated in fission yeast need to be revisited using FXm. Together with the wealth of knowledge on size control in this organism, this approach will provide new insights into the processes that underlie the biology of cell growth. Interestingly, the investigation of biological processes has been enriched by the use of natural yeast isolates. In fission yeast, these strains are known to be less homogenous in morphology and overall size (Jeffares et al., 2015). FXm may therefore make it possible to fully exploit this unique resource and take advantage of advanced population genetics strategies to decipher the regulation of cell size. While we use fission yeast as a proof-of-concept model, our results also show that FXm can be applied to any type of cell over a broad range of sizes. Our studies therefore establish the versatility and power of FXm for investigating cell volume homeostasis and its regulation in small cells and model organisms, contributing to our general understanding of the modulation of cellular dimensions and scaling in eukaryotes.

## MATERIALS AND METHODS

### Fission yeast strains and methods

Standard methods and media were used (Hayles and Nurse, 1992; Moreno et al., 1991). All the strains described in this study are detailed in Table S2. The deletions of *rga2*, *rga4*, *rga6*, *mal3*, *knk1* and *tea1* have already been described (Soto et al., 2010; Villar-Tajadura et al., 2008) (Beinhauer et al., 1997; Mata and Nurse, 1997; Revilla-Guarinos et al., 2016; Scheffler et al., 2014). All experiments were carried out in non-supplemented minimal medium (EMM) at 32 °C, except otherwise noted. To inhibit CDK activity in analog-sensitive MCN strains, the 3-MBPP1 inhibitor (A602960, Toronto Research Chemicals, Inc.) was dissolved in DMSO at stock concentrations of 10 or 0.4 mM and added to liquid cultures at a final concentration of 1 or 0.05 μM. The percentage of binucleated cells (Fig. S3C) was determined from heat fixed samples on microscope slides (70 °C for 5 min) stained with a 11:1 solution of Blankophor (1 mg/mL): DAPI (1 μg/mL).

### Microfabrication of FXm chips

PDMS microfluidic chips for FXm were prepared following standard microfabrication protocols (McDonald and Whitesides, 2002). In brief, master molds were made by spin-coating SU-8 2005 resin (MicroChem Corp., USA) on silicon wafers using a spincoater (Laurell Technologies, USA) according to manufacturer’s instructions. Microstructures were then generated using high-resolution chrome masks (JD phototools, UK) and 365nm UV exposure (UV KUB 3, Kloe, France) followed by PGMEA (Sigma-Aldrich) development. For each mold, the height of the structures was determined as the average of three measurements perpendicular to the long axis of the design (Fig. 1B) using a Veeco Wyko NT9100 optical profilometer (Veeco Instruments Inc., USA). Note that molds showing significant variability between the three height measurements should be either discarded or used considering local rather than global averaged height. To produce chips for FXm, a 10:1 mixture of PDMS (Sylgard 184, Dow Corning, USA) was cast on the SU-8 master mold and allowed to cure at 70 °C for 2 hours. Inlets were then made using 2.5 or 4 mm biopsy punches, and chips were bonded to microscopy-grade coverslips by plasma activation (Harrick Plasma, USA).

### Preparation of fission yeast cells for FXm

To perform FXm experiments, exponentially growing fission yeast cells were sampled at an optical density between 0.25 and 0.45 OD_595_. To limit the formation of cell aggregates, a mild sonication cycle of 5 s at 10 % amplitude (Branson 450 Digital Sonifier, Emerson Electric Co.) was applied. This treatment had no effect on the results we obtained by FXm (data not shown). In order to optimize the number of cells that can be measured on a single FXm image in our chip design, cells were concentrated to ~5.5×10^7^ cells/mL in their own conditioned minimal medium (note that to prevent bias in our volume measurements, we did not use autofluorescent rich medium). FITC-Dextran was then added to the sample (FD10S, Sigma Aldrich) at a final concentration of 1 mg/mL, and cells were loaded into the chip shortly before image acquisition. For evaluating the volume of quiescent cells, cells were sampled 24 hours after they reached 0.4 OD_595_.

### Microscopy

All microscopy experiments were carried out using an inverted Zeiss Axio Observer (Carl Zeiss Microscopy LLC) equipped with a Lumencor Spectra X illumination system and an Orca Flash 4.0V2 sCMOS camera (Hamamatsu Photonics). Acquisition was performed using the VisiView software (Visitron Systems GmbH). A Plan-Apochromat 63X/1.4 NA immersion lens and a Plan-Apochromat 20X/0.8 NA Ph2 lens (Carl Zeiss Microscopy LLC) were used for cell length measurements and FXm, respectively. For routine volume measurements, the acquisition parameters were 100 ms exposure at 20 % illumination power, and 30 to 40 images were taken across the whole chambers to limit potential bias due to local changes in chamber height. For the time-lapse experiments in Fig 8, 100 ms exposure at 10 % illumination power was used.

### Image analysis

Cell length at division was determined from Blankophor images (1 mg/mL Blankophor solution, A108741, Ambeed Inc.) using FIJI (National Institutes of Health) and the Pointpicker plug-in. For the blankophor images in Fig. 2, 4, 5 and 7, the brightness and contrast were adjusted for display purposes. For routine volume measurements, images were first normalized using a custom Matlab software (Cadart et al., 2017). Cell selection and volume calculation were subsequently performed using the normalization mask with a new, specifically-developed Python interface (see Results). For all Figures, cell selection was done using the manual mode, except for Fig. 3B and 4C where automated cell identification was used. Note that except for the time-lapses in Fig. 8B and C, we systematically excluded cells that were undergoing cytokinesis but that had not fully separated into two individual daughter cells; this is similar to studies of cell length at division. While this category can be integrated as newly-born pairs of cells using the original Matlab software (Cadart et al., 2017), manually delineating the division site would introduce measurement errors. Finally, our Python code is not adapted to tracking the volume of individual cells in long time-lapse experiments (Fig. 8B, C), as slight cell movements make it difficult to reliably follow single cells over time when using the normalization mask pre-selection. For these assays, we used the original Matlab software (Cadart et al., 2017).

### FXm time-lapse and analysis using fission yeast

As fission yeast cells are non-adherent, FXm time-lapses required coverslips to be coated in order to prevent cells from moving inside the chamber, which would hamper data acquisition and analysis. To this end, 10 μl of 1mg/ml filtered lectin was spread in a rectangle of ~10×5 mm at the center of the coverslip and allowed to dry at 37 °C. The coverslip was then washed once with filtered ultra-pure water and dried at 37 °C. For experiments without medium flow (Fig. 8B), Ø 4 mm inlets were made in the PDMS chip using the appropriate biopsy punch. 5 μl of cells prepared as described above were loaded in one inlet, and both inlets were then filled with 45 μl EMM containing the FITC-Dextran dye. Cells were allowed to flow into the chamber prior to imaging. For time-lapse experiments using medium flow (Fig. 8C), two successive layers of lectin were applied as above and inlets of Ø 0.75 mm were fabricated. Cells were then loaded by depositing a drop of cells prepared as above over one of the inlets, and a mild vacuum was applied at the other inlet using the tip of a 10 mL pipet connected to a vacuum pump. The loaded chip was then connected to a flow control system (pressure generator and flow controller, Elvesys, France), and a flow of medium (~3-5 μL/ min) containing the FITC-Dextran dye was used throughout the experiment. For the analysis of the time-lapse experiments, the two daughter cells after a division were manually delineated during the analysis and only one of them was followed for further investigation (Fig. 8B, C). The growth rate of each individual cell was determined by calculating the slopes of the traces during each cycle. This was obtained by linear regression considering the volume at birth as the first time point (when an invagination is detected) and excluding the mitotic plateau: entry into mitosis was assigned to the last point prior to strong reduction of growth (when two consecutive points on the graph decrease the slope).

## Supporting information

Supplemental code for FXm analysis

## ACKNOWLEDGMENTS

We thank Pei-Yun Jenny Wu and Snezhana Oliferenko for critically reading the manuscript. We thank Pascal Hersen and Giacomo Gropplero for the measurements of our chamber heights. We are grateful to Paul Nurse, Snezhana Oliferenko, Pilar Perez and Phong Tran for sharing strains and reagents.

