## Supplemental Information for "Fluorescence exclusion: a rapid, accurate and powerful method for measuring yeast cell volume"

### Figure S1

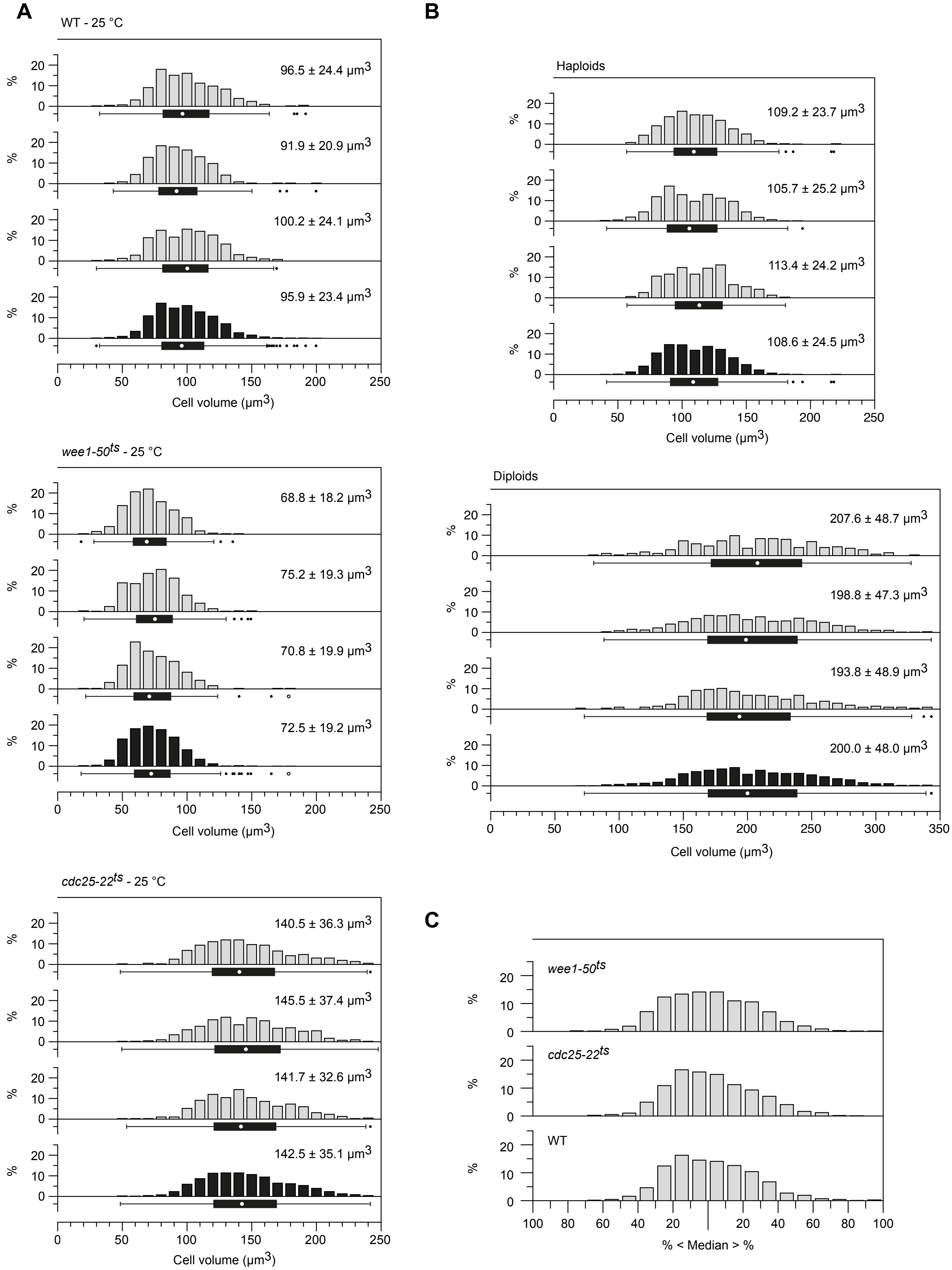

A
WT - 25 °C
96.5 ± 24.4 µm3
91.9 ± 20.9 µm3
100.2 ± 24.1 µm3
95.9 ± 23.4 µm3
Cell volume (µm3)
wee1-50ts - 25 °C
68.8 ± 18.2 µm3
75.2 ± 19.3 µm3
70.8 ± 19.9 µm3
72.5 ± 19.2 µm3
Cell volume (µm3)
cdc25-22ts - 25 °C
140.5 ± 36.3 µm3
145.5 ± 37.4 µm3
141.7 ± 32.6 µm3
142.5 ± 35.1 µm3
Cell volume (µm3)
B
Haploids
109.2 ± 23.7 µm3
105.7 ± 25.2 µm3
113.4 ± 24.2 µm3
108.6 ± 24.5 µm3
Cell volume (µm3)
Diploids
207.6 ± 48.7 µm3
198.8 ± 47.3 µm3
193.8 ± 48.9 µm3
200.0 ± 48.0 µm3
Cell volume (µm3)
C
wee1-50ts
cdc25-22ts
WT
% < Median > %

**Figure S1. Supporting data for Fig. 2. A, B.** Volume measurement by FXm of the indicated strains. All the replicates (grey) used for the data presented in Fig. 2C (panel *A*) and Fig. 2D (panel *B*) are shown. For each strain, the bottom histogram (black) and box plot are the same as in Fig. 2C, D. Graphs are as in Fig. 2. For each replicate in *A*: 1) wild type (WT): $n \geq 337$, *cdc25-22*$^{ts}$: $n \geq 353$, *wee1-50*$^{ts}$: $n \geq 374$ and 2) values outside of the plot range were excluded (WT: $n^*=0$; *cdc25-22*$^{ts}$: $n^* \leq 4$, *wee1-50*$^{ts}$: $n^*=0$). For each replicate in *B*: 1) Haploids: $n \geq 311$, diploids: $n \geq 207$ and 2) values outside of the plot range were excluded (haploids: $n^*=0$; diploids: $n^* \leq 1$). **C.** Volume distribution for the indicated strains as a percentage of the median volume of the population. Datasets as in Fig. 2C and Fig. S1A. Values outside of the plot range were excluded ($n^* \leq 4$).

### Figure S2

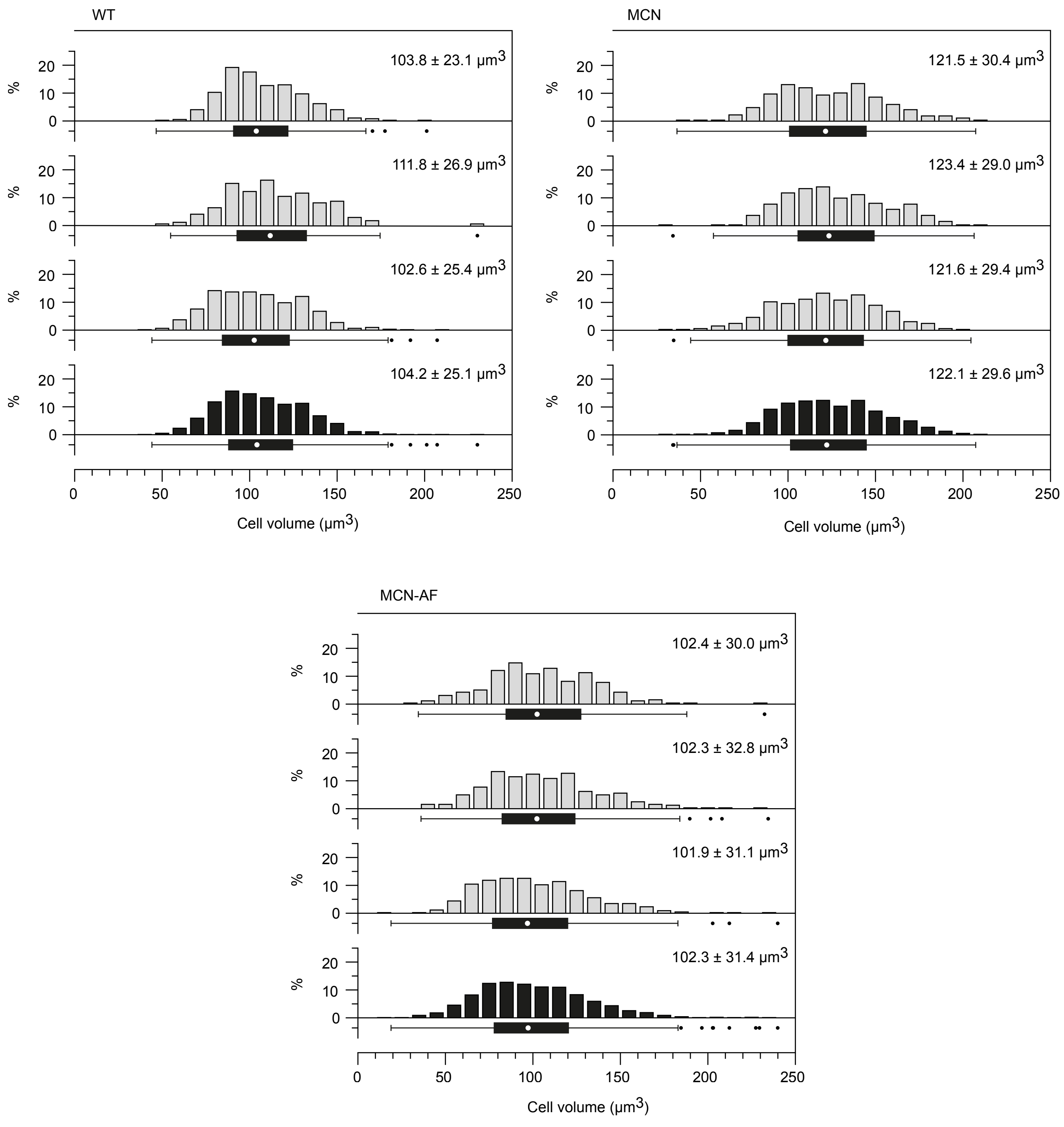
WT
MCN
MCN-AF
103.8 ± 23.1 μm3
111.8 ± 26.9 μm3
102.6 ± 25.4 μm3
104.2 ± 25.1 μm3
121.5 ± 30.4 μm3
123.4 ± 29.0 μm3
121.6 ± 29.4 μm3
122.1 ± 29.6 μm3
102.4 ± 30.0 μm3
102.3 ± 32.8 μm3
101.9 ± 31.1 μm3
102.3 ± 31.4 μm3
%
Cell volume (μm3)

**Figure S2. Supporting data for Fig. 4A.** Volume measurement by FXm of the indicated strains. All replicates (grey) used for the data presented in Fig. 4A are shown. For each strain, the bottom histogram (black) and box plot are the same as in Fig. 4A. For wild type (WT), the 1st and 3rd datasets are as in Fig. 2A (last and 1st datasets respectively). Graphs are as in Fig. 2. For each replicate: 1) WT: $n \geq 172$, MCN: $n \geq 267$, MCN-AF: $n \geq 257$ and 2) values outside of the plot range were excluded (WT: $n^* = 0$; MCN: $n^* = 0$, MCN-AF: $n^* \leq 1$).

#### Figure S3

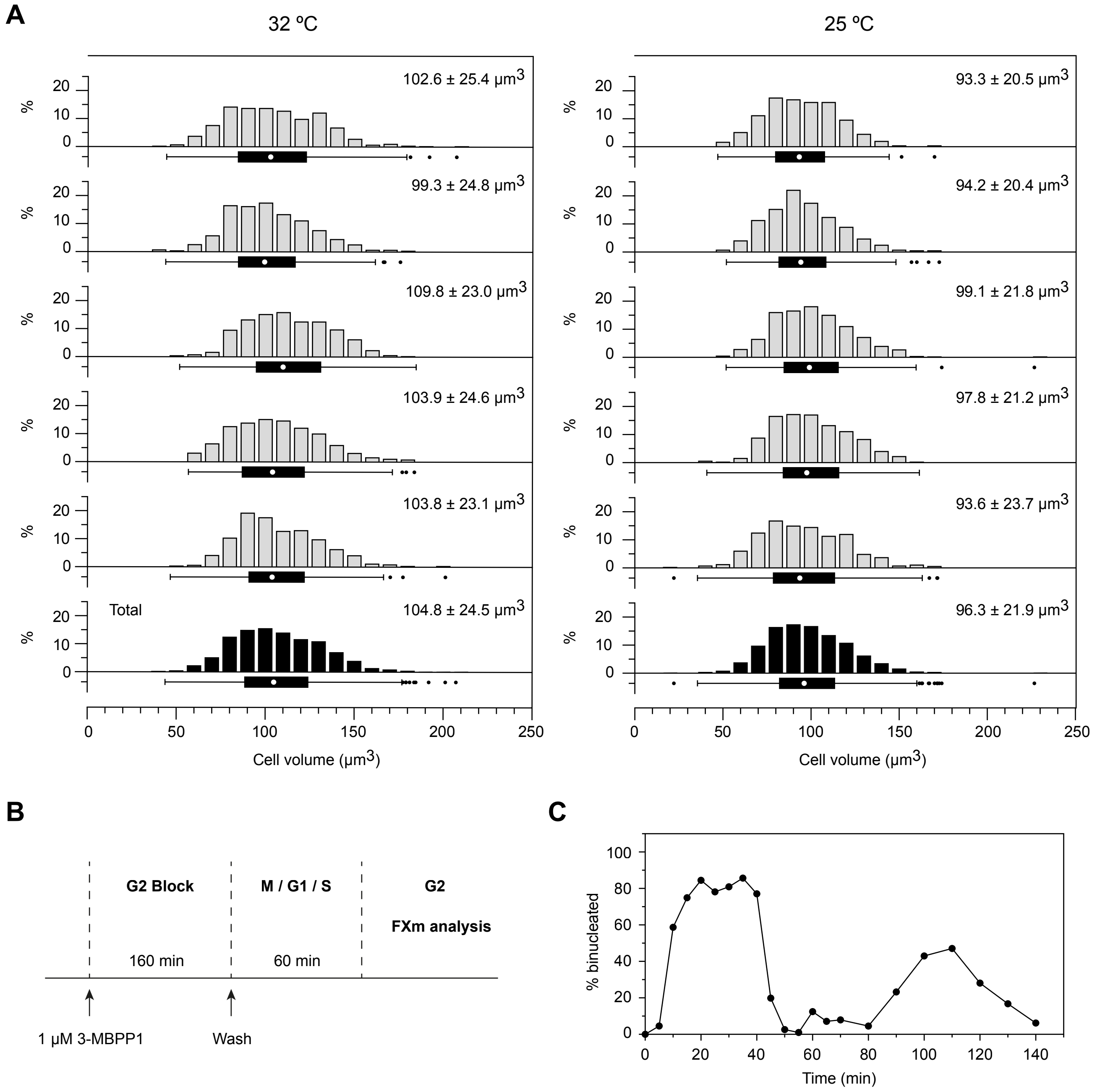

A
32 ºC
25 ºC
102.6 ± 25.4 µm3
99.3 ± 24.8 µm3
109.8 ± 23.0 µm3
103.9 ± 24.6 µm3
103.8 ± 23.1 µm3
Total
104.8 ± 24.5 µm3
93.3 ± 20.5 µm3
94.2 ± 20.4 µm3
99.1 ± 21.8 µm3
97.8 ± 21.2 µm3
93.6 ± 23.7 µm3
96.3 ± 21.9 µm3
%
Cell volume (µm3)
B
G2 Block
M / G1 / S
G2
FXm analysis
160 min
60 min
1 µM 3-MBPP1
Wash
C
% binucleated
Time (min)

**Figure S3. Supporting data for Fig. 4B and 4C. A.** Measurements for five independent experiments (grey) and for the total pooled datasets (black) using wild type cells grown at 32 °C (left panel) or 25 °C (right panel). Data at 32 °C are as in Fig. 2A. For the data at 25 °C, $n \geq 316$ for each replicate, $n=2706$ for the total dataset; no values were outside of the plot range ($n^*=0$ for each replicate). Graphs are as in Fig. 2. The median volume values with standard deviations are also shown in Fig. 4B. **B.** Schematic of the experimental protocol for the data in Fig. 4C. **C.** % of binucleated cells after synchronization ($n \geq$ 178 at each time point). Samples were analyzed at 5 min intervals for the first 70 min and then at 10 min intervals. This experiment is independent from that in Fig. 4C and was entirely performed in culture flasks. It was used to determine the time window after the release from the G2 block during which images would be acquired for Fig. 4C.

**Figure S4**

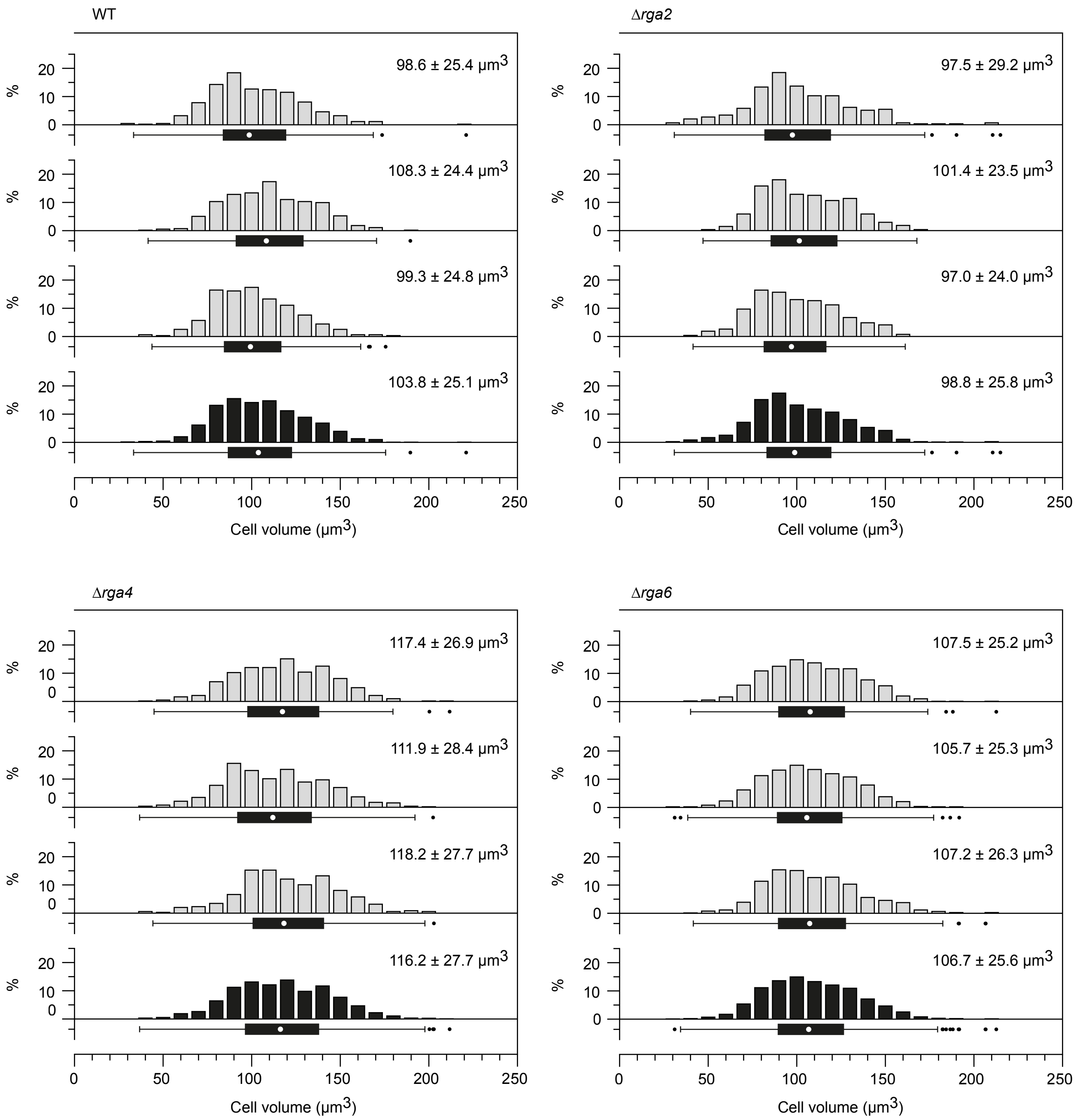
WT
98.6 ± 25.4 µm3
108.3 ± 24.4 µm3
99.3 ± 24.8 µm3
103.8 ± 25.1 µm3
Δrga2
97.5 ± 29.2 µm3
101.4 ± 23.5 µm3
97.0 ± 24.0 µm3
98.8 ± 25.8 µm3
Δrga4
117.4 ± 26.9 µm3
111.9 ± 28.4 µm3
118.2 ± 27.7 µm3
116.2 ± 27.7 µm3
Δrga6
107.5 ± 25.2 µm3
105.7 ± 25.3 µm3
107.2 ± 26.3 µm3
106.7 ± 25.6 µm3
%
0
10
20
0
50
100
150
200
250
Cell volume (µm3)

**Figure S4. Supporting data for Fig. 5B.** Volume measurement by FXm of the indicated strains. All replicates (grey) used for the data presented in Fig. 5B are shown. For each strain, the bottom histogram (black) and box plot are the same as in Fig. 5B. For wild type (WT), the $1^{st}$ dataset is as in Fig. 2A ($4^{th}$ dataset). Graphs are as in Fig. 2. For each replicate: 1) WT: $n \geq 316$, Δ*rga2*: $n \geq 268$, Δ*rga4*: $n \geq 348$, Δ*rga6*: $n \geq 766$ and 2) values outside of the plot range were excluded (WT: $n^* \leq 1$; Δ*rga2*: $n^*=0$, Δ*rga4*: $n^*=0$, Δ*rga6*: $n^*=0$).

### Figure S5

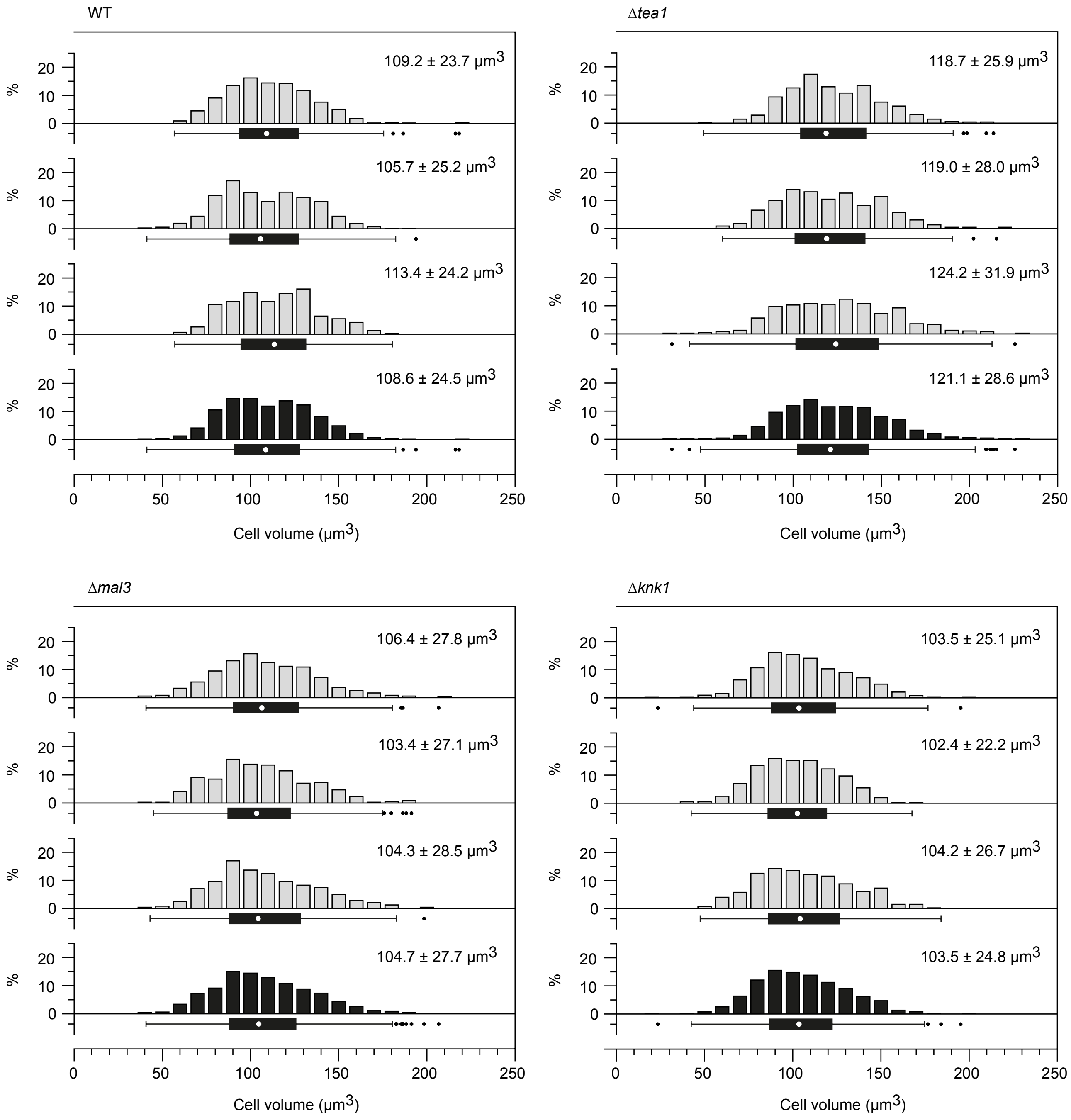

WT
109.2 ± 23.7 μm3
105.7 ± 25.2 μm3
113.4 ± 24.2 μm3
108.6 ± 24.5 μm3
Δtea1
118.7 ± 25.9 μm3
119.0 ± 28.0 μm3
124.2 ± 31.9 μm3
121.1 ± 28.6 μm3
Δmal3
106.4 ± 27.8 μm3
103.4 ± 27.1 μm3
104.3 ± 28.5 μm3
104.7 ± 27.7 μm3
Δknk1
103.5 ± 25.1 μm3
102.4 ± 22.2 μm3
104.2 ± 26.7 μm3
103.5 ± 24.8 μm3
%
Cell volume (μm3)

**Figure S5. Supporting data for Fig. 5D.** Volume measurement by FXm of the indicated strains. All the replicates (grey) used for the data presented in Fig. 5D are shown. For each strain, the bottom histogram (black) and box plot are the same as in Fig. 5D. Graphs are as in Fig. 2. For each replicate: 1) wild type (WT): $n \geq 311$, Δ*tea1*: $n \geq 230$, Δ*mal3*: $n \geq 242$, Δ*knk1*: $n \geq 398$ and 2) no values were outside of the plot range ($n^*=0$ for each strain).

**Figure S6**

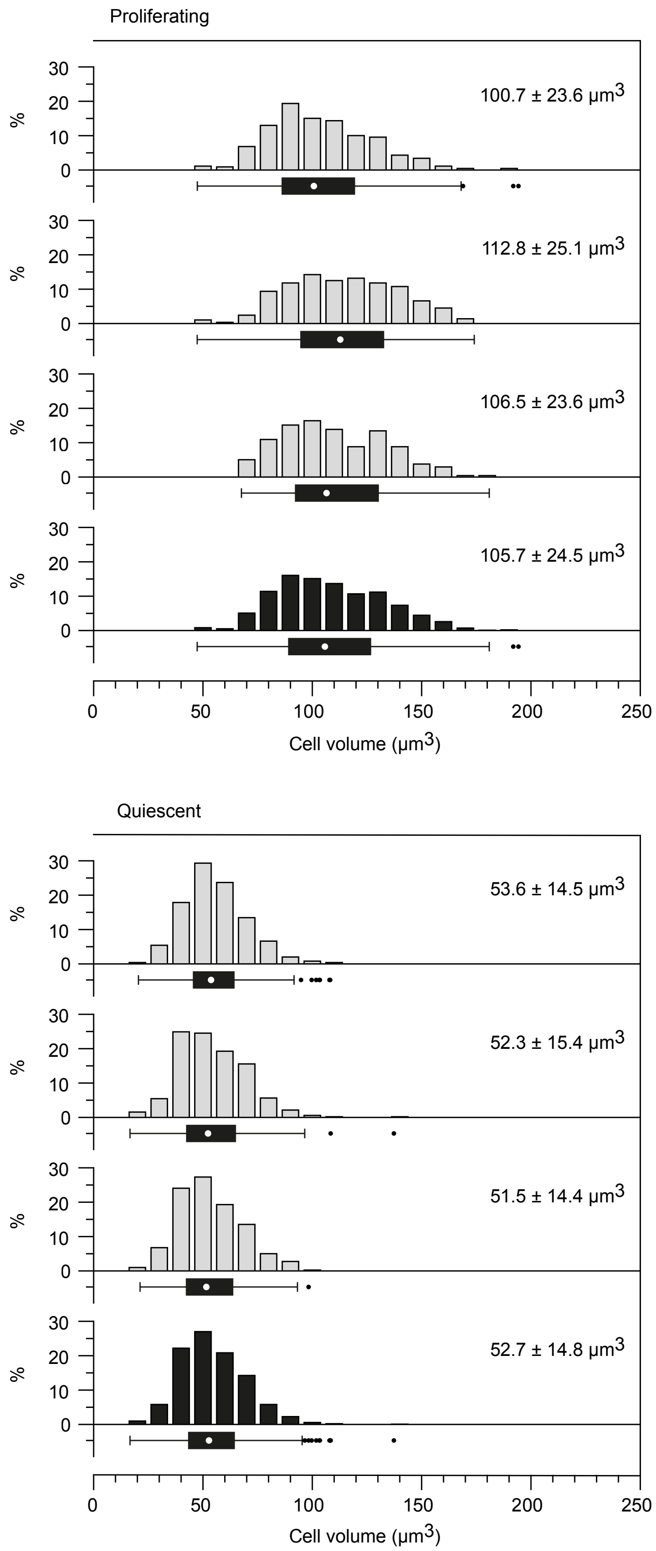

Proliferating
100.7 ± 23.6 µm3
112.8 ± 25.1 µm3
106.5 ± 23.6 µm3
105.7 ± 24.5 µm3
%
0
10
20
30
0
50
100
150
200
250
Cell volume (µm3)
Quiescent
53.6 ± 14.5 µm3
52.3 ± 15.4 µm3
51.5 ± 14.4 µm3
52.7 ± 14.8 µm3
Cell volume (µm3)

**Figure S6. Supporting data for Fig. 5F.** Volume measurement by FXm of proliferating *vs*. quiescent wild type fission yeast cells. All the replicates (grey) used for the data presented in Fig. 5F are shown. For each strain, the bottom histogram (black) and box plot are the same as in Fig. 5F. Graphs are as in Fig. 2. For each replicate: 1) proliferating: $n \geq 238$, quiescent: $n \geq 399$ and 2) no values were outside of the plot range ($n^*=0$).

### Figure S7

**A**

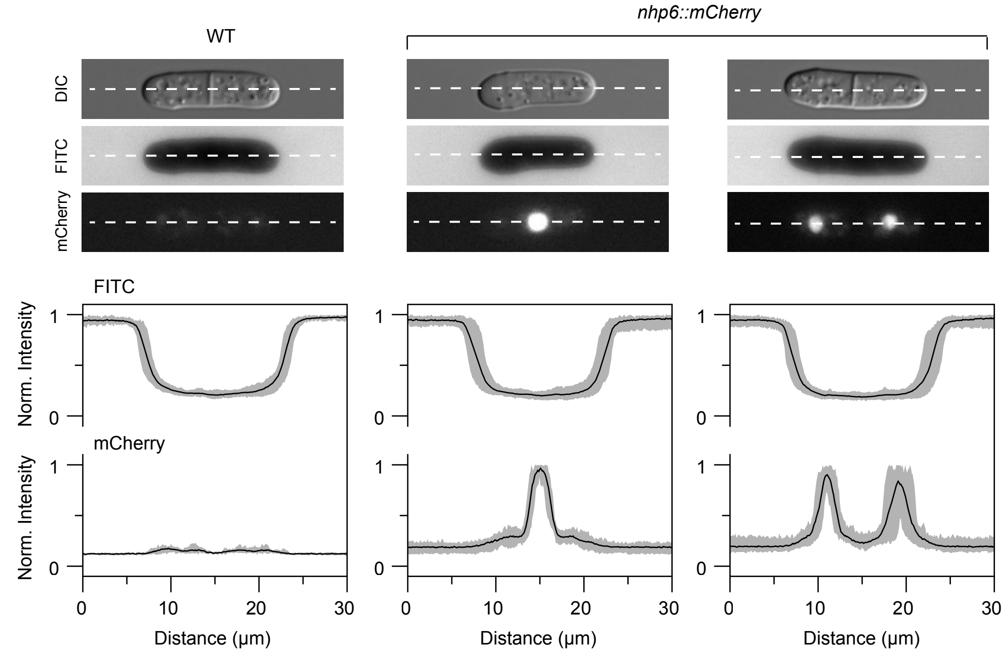

WT
nhp6::mCherry
DIC
FITC
mCherry
FITC
Norm. Intensity
mCherry
Norm. Intensity
1
0
0
10
20
30
Distance (μm)


**B**

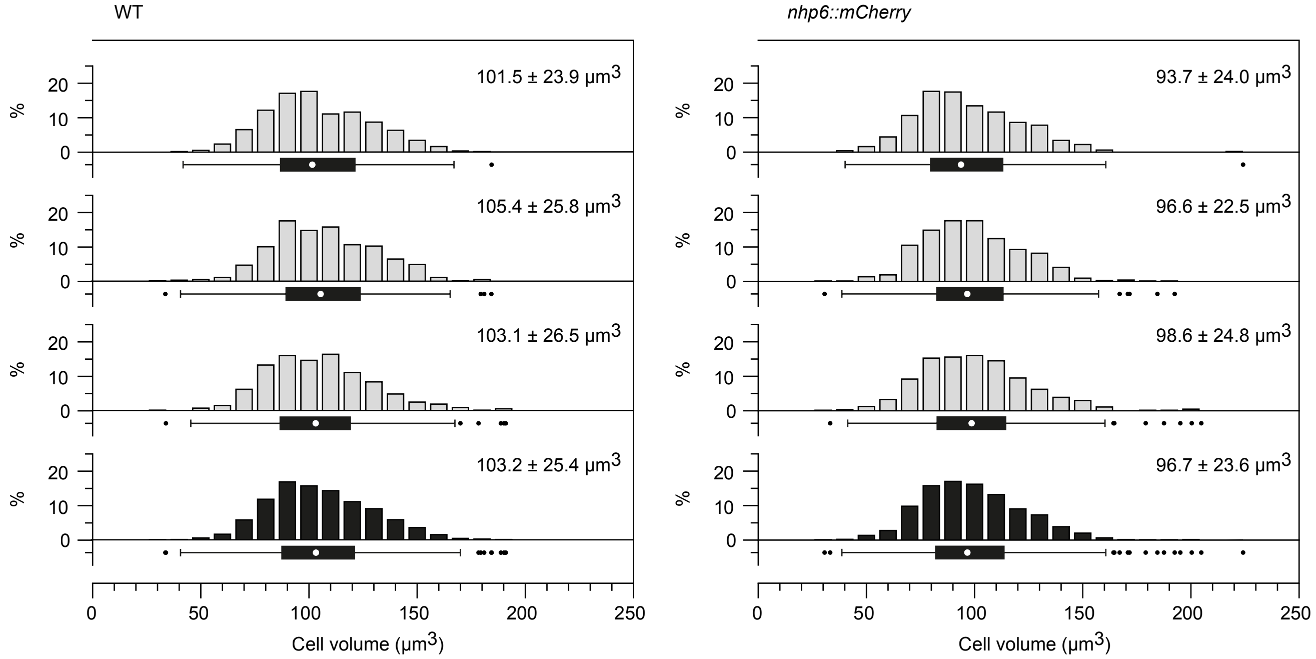

WT
nhp6::mCherry
101.5 ± 23.9 μm3
105.4 ± 25.8 μm3
103.1 ± 26.5 μm3
103.2 ± 25.4 μm3
93.7 ± 24.0 μm3
96.6 ± 22.5 μm3
98.6 ± 24.8 μm3
96.7 ± 23.6 μm3
%
0
10
20
50
100
150
200
250
Cell volume (μm3)

**Figure S7. Supporting data for Fig. 6. A.** Top panel: representative images of wild type (WT) and *nhp6::mCherry* cells in the DIC (top), FITC (middle) and mCherry (bottom) channels. Equivalent numbers of cells of both strains were mixed and imaged in the presence of the FITC-Dextran dye. Dashed lines indicate the path of the fluorescence intensity scans presented in the bottom panel. Bottom panel: quantification of the fluorescence intensity in both channels. 30 µm scan lines were drawn using Fiji (National Institutes of Health), manually centered on the cells, and fluorescence intensities along the scan lines were determined. In the FITC channel as well as in the mCherry channel for *nhp6::mCherry* cells, values were normalized to the maximum intensities measured for each cell. For wild type in the mCherry channel, values were normalized to the maximum intensity measured in the complete dataset of Nhp6::mCherry positive cells. Black lines: averages of all the intensity profiles ($n$=12 for each cell type). The grey areas are delineated by the maximum and minimum intensities at each position in the complete datasets. **B.** Volume measurement by FXm of the indicated strains. All the replicates (grey) used for the data presented in Fig. 6A are shown. For each strain, the bottom histogram (black) and box plot are the same as in Fig. 6A. Graphs are as in Fig. 2. For each replicate: 1) WT: $n \geq 506$, *nhp6::mCherry*: $n \geq 499$ and 2) values outside of the plot range were excluded (WT: $n^* \leq 1$; *nhp6::mCherry*: $n^*=0$).

**Table S1. Comparison of strains based on volume *vs.* length at division**

| | | Comparison | V* ratio | SAD** ratio | Data |
|---|---|---|---|---|---|
| *cdc25-22*$^{ts}$ at 25 ºC | | WT at 25 ºC | 1.49 | 1.36 | Fig. 2C |
| *wee1-50*$^{ts}$ at 25 ºC | | WT at 25 ºC | 0.76 | 0.79 | Fig. 2C |
| WT diploids in EMM6S | | WT haploids in EMM6S | 1.84 | 1.44 | Fig. 2D |
| MCN | | WT | 1.17 | 1.07 | Fig. 4A |
| MCN-AF | | WT | 0.98 | 0.91 | Fig. 4A |
| MCN-AF | | MCN | 0.84 | 0.85 | Fig. 4A |
| Δ*rga2* | | WT | 0.95 | 1.17 | Fig. 5A, B |
| Δ*rga4* | | WT | 1.12 | 0.88 | Fig. 5A, B |
| Δ*rga6* | | WT | 1.03 | 0.96 | Fig. 5A, B |
| *nhp6::mCherry* | | WT | 0.94 | 0.98 | Fig. 6A |
| MCN + 3-MBPP1 | 40 min | MCN + 3-MBPP1 / 0 min | 1.07 | 1.05 | Fig. 7 |
| | 80 min | MCN + 3-MBPP1 / 0 min | 1.27 | 1.39 | Fig. 7 |
| | 120 min | MCN + 3-MBPP1 / 0 min | 1.44 | 1.55 | Fig. 7 |
| | 160 min | MCN + 3-MBPP1 / 0 min | 1.72 | 1.58 | Fig. 7 |
| WT at 25 ºC | | WT at 32 ºC | 0.92 | 0.93 | Fig. 2C, 4A |
| WT in EMM6S at 32 ºC | | WT in EMM at 32 ºC | 1.04 | 0.93 | Fig. 2D, 4A |

* V: volume; ** SAD: cell size at division

**Table S2. Fission yeast strains used in this study**

| **Name** | **Genotype** | **Origin** |
|---|---|---|
| PN1 | *h- 972* | P. Nurse |
| DC1114 | *h-/h- 972 diploid* | This study |
| PN143 | *h- cdc25-22* | P. Nurse |
| PN369 | *h- wee1-50* | P. Nurse |
| DC676 | *h- Pcdc13::cdc13Scdc2as::cdc13UTR Δcdc2::KAN Δcig1::HYG Δcig2::KAN Δpuc1::HYG* | This study |
| DC753 | *h- Pcdc13::cdc13Scdc2T14AY15Fas::cdc13UTR Δcdc2::KAN Δcig1::HYG Δcig2::KAN Δpuc1::HYG* | This study |
| DC1099 | *h+ Δrga2::KAN* | This study |
| DC1102 | *h- Δrga4::KAN* | This study |
| DC1100 | *h+ Δrga6::KAN* | This study |
| DC1131 | *h- nhp6::mCherry::ura4+* | This study |
| PN1687 | *h- Δtea1::ura4+ ura4-D18* | P. Nurse |
| PN3718 | *h- Δmal3::his3 ade6-M210 leu1-32 ura4-D18 his3-D1* | P. Nurse |
| TP474 | *h- Δknk1::KAN ade6-M210 leu1-32 ura4-D18* | P. Tran |
