## Supplemental code for FXm analysis for "Fluorescence exclusion: a rapid, accurate and powerful method for measuring yeast cell volume": autoSegment.py

```
import numpy as np
import pandas as pd
import mahotas as mh


def get_bg_box(x, y, img_size=2048):
    """
    Calculates coordinates of a 200x200 box without overlapping with the edges
    :param x: x coordinate of center
    :param y: y coordinate of center
    :param img_size: size of the image, used to clip the box coordinates to the borders
    :return coords: list of coordinates for the 200x200 box
    """
    # Calculates coordinates of 200x200 box
    coords = [x - 100, x + 100, y - 100, y + 100]
    for i in range(len(coords)):
        if coords[i] < 0:
            coords[i] = 0
        elif coords[i] > img_size:
            coords[i] = img_size
    return coords


def segment(pillar_mask):
    """
    Filters regions drawn by the normalization script in pillar_mask
    :param pillar_mask: mask with background = False and cells and pillars = True.
    :return cells: image with labeled regions
    """

    # Uses pillar_mask to find separated cells
    pillar_mask, n_elem = mh.label(pillar_mask)
    sizes = mh.labeled.labeled_size(pillar_mask)
    pillar_mask = mh.labeled.remove_regions(pillar_mask, np.where(sizes > 20000))  # Threshold size to remove pillars
    pillar_mask = mh.labeled.remove_bordering(pillar_mask)  # Removes selections touching the edges
    cells, n_cells = mh.labeled.relabel(pillar_mask)

    return cells


def get_volume(img, bg_mask, cells, pillar_height=5.6, pixel_size=0.325):
    """
    Calculates cell volume based on input parameters
    :param img: image
    :param bg_mask: mask with background = 0 and cells and pillars = 127.
    :param cells: labeled image with cell selections
    :param pillar_height: Height of microfluidic chamber
    :param pixel_size: Size of pixel given camera pixel size and microscope magnification
    :return df: DataFrame with all image analysis parameters, including cell volume
    """

    img_size = img.shape[0]

    # Gets values of intensity, surface and center of mass
    surfaces = mh.labeled.labeled_size(cells)
    centers = mh.center_of_mass(cells, cells)

    # Removes background region from data
    surfaces = surfaces[1:]
    centers = centers[1:]

    intensities = []
    bg_intensities = []
    volumes = []
    for c, pix in enumerate(centers):  # Iterates over all intensity centers
        x = int(pix[0])
        y = int(pix[1])

        [x0, x1, y0, y1] = get_bg_box(x, y, img_size=img_size)

        # Gets mask corresponding to region, applies it and calculates mean intensity (darkness of cells)
        current_mask = (cells == c + 1)
        img_masked = img[current_mask]
        mean = np.mean(img_masked)
        intensities.append(mean)

        # Creates 200x200 box in both image and background and calculates median intensity (brightness of background)
        selection = img[x0:x1, y0:y1]
        selected_bg = bg_mask[x0:x1, y0:y1]
        median = np.median(selection[np.where(selected_bg == 0)])

        """
        Basis of the FXm:
        median - mean = difference between background intensity and cell intensity (excluded fluorescence)
        surface = number of pixels
        pillar_height = height of the microfluidic chamber (µm)
        pixel_size = camera pixel size (µm)
        """
        vol = (median-mean) * surfaces[c] * pillar_height * pixel_size ** 2
        volumes.append(vol)

        bg_intensities.append(median)

    df = pd.DataFrame({"Path": None,
                       "ID": list(range(len(centers))),
                       "Center X": list(centers[:, 0]),
                       "Center Y": list(centers[:, 1]),
                       "Pillar Height": [pillar_height] * len(centers),
                       "Pixel Size": [pixel_size] * len(centers),
                       "Background": list(bg_intensities),
                       "Surface": list(surfaces),
                       "Intensity": list(intensities),
                       "Volume": list(volumes)})

    return df
```
