## Supplemental code for FXm analysis for "Fluorescence exclusion: a rapid, accurate and powerful method for measuring yeast cell volume": group.py

```
import numpy as np
import matplotlib.pyplot as plt
from matplotlib.widgets import RadioButtons
import autoSegment as auto
import mahotas as mh
import pandas as pd
import os.path
import argparse


def on_click(event):
    """
    Executed when user clicks on an object.
    Changes group assigned to object and displays it accordingly.
    """
    global ix
    ix = event.xdata
    axes = event.inaxes

    if not axes or not axes in ax:  # Catches when user does not click on an image
        return

    # Finds clicked axes
    index = ax.index(axes)

    # Group handling
    palette = ["lime", "yellow", "cyan", "darkorange", "skyblue", "salmon", "blueviolet", "lightgray"]
    global ng
    old_group = preselection[index]
    
    if event.button == 1:
        new_group = (old_group + 1) % (ng + 1)
    elif event.button == 3:
        new_group = (old_group - 1) % (ng + 1)

    # Formats group labels
    if new_group == 0:
        group_label = "None"
        group_color = "red"

    else:
        group_label = f"G{new_group}"
        group_color = palette[new_group % len(palette) - 1]

    axes.texts[-1].remove()
    axes.text(10, 100, group_label, bbox=dict(facecolor=group_color, alpha=1))
    plt.draw()

    preselection[index] = new_group


def filter_cells(img, marker, mask, cells_df, path=None, ng=0):
    """
    Displays segmented cells and allows the user to assign them to different groups
    :param img: original image
    :param marker: image of the marker used to differentiate groups
    :param mask: original segmentation mask
    :param cells_df: DataFrame with volume data
    :param path: path of the image being analyzed. Will be displayed as the window title.
    :param ng: Number of groups to classify the cells in
    :return df: DataFrame with updated "Filtered" column (True for discarded objects)
    """
    print(path)

    img_size = img.shape[0]

    # Initializes bool list with groups.
    global preselection
    if "ManualFilter" in cells_df.head():
        # Analysis file from manual mode: Only shows objects accepted as cells
        preselection = list(cells_df["ManualFilter"] == False)
        if sum(preselection) == 0:
            # There are no cells
            preselection = [0] * len(cells_df["Volume"])
            print("There are no cells in this image")
            return preselection

        else:
            n_cells = sum(preselection)  # Number of cells to show (+1 for the radiobutton)

    else:
        # Analysis file from automatic mode: Shows all objects
        n_cells = len(cells_df["Volume"]) # Number of cells to show (all, +1 for the radiobutton)
        preselection = [True] * n_cells

    # Plotting parameters
    rows = np.ceil(np.sqrt(n_cells + 1))
    columns = rows
    fig = plt.figure(figsize=(16, 9))  # Adapts these parameters for smaller screens
    fig.suptitle(path)

    centers = cells_df[["Center X", "Center Y"]].to_numpy()

    # Saves subplots in a list to be able to access them
    global ax, fxm_list, marker_list
    ax = []
    fxm_list = []
    marker_list = []

    j = 0  # Counts objects that have been shown
    for i, pix in enumerate(centers):
        # Checks if item should be displayed
        if not preselection[i]:
            preselection[i] = 0  # Sets group to 0
            ax.append(None)
            continue

        preselection[i] = 1  # Sets here the initial group

        # Adds new empty plot to selection window
        subp = fig.add_subplot(rows, columns, j + 1)

        # Formatting options
        subp.set_yticklabels([])
        subp.set_xticklabels([])
        ax.append(subp)

        # Creates group label
        ax[-1].text(10, 100, 'G1', bbox=dict(facecolor='lime', alpha=1))

        # Gets center coordinates of object
        x = int(pix[0])
        y = int(pix[1])

        # Gets 200x200 pixels area around object
        [x0, x1, y0, y1] = auto.get_bg_box(x, y, img_size=img_size)
        msk_img = img[x0:x1, y0:y1] * (mask[x0:x1, y0:y1] > 0)

        fxm_img = plt.imshow(msk_img)
        fxm_list.append(fxm_img)
        msk_marker = marker[x0:x1, y0:y1] * (mask[x0:x1, y0:y1] > 0)

        marker_img = plt.imshow(msk_marker, cmap=plt.cm.Reds, alpha=1)  # Changes the color map if your marker is not red
        marker_list.append(marker_img)

        # Uncomments next line to draw point to identify cell
        # plt.scatter(100, 100, s=1, color="red")

        j += 1  # Update counter of displayed objects

    cid = fig.canvas.mpl_connect('button_press_event', on_click)  # Listens for click on object

    def change_channel(label):
        global ax, fxm_list, marker_list
        if label == "FXm":
            for c, v in enumerate(fxm_list):
                v.set_alpha(1)
                marker_list[c].set_alpha(0)

        elif label == "Marker":
            for c, v in enumerate(fxm_list):
                v.set_alpha(0)
                marker_list[c].set_alpha(1)

        plt.draw()

    def on_press(event):
        if event.key == ' ':
            sel = radio.value_selected
            for c, v in enumerate(radio.labels):
                # Finds the non-selected label
                label = v.get_text()
                if sel != label:
                    radio.set_active(c)
                    break

    # Creates new subplot with the radiobuttons
    rax = fig.add_subplot(rows, columns, j + 1)
    radio = RadioButtons(rax, ("Marker", "FXm"))
    radio.on_clicked(change_channel)

    fig.canvas.mpl_connect('key_press_event', on_press)

    plt.show()

    return preselection


# User interaction and parameter logic
parser = argparse.ArgumentParser(description="Separate FXm cells in groups")
parser.add_argument("path", type=str, help="Path to .tsv analysis file")
parser.add_argument("-g", "--groups", type=int, help="Number of groups to classify the cells in.")
parser.add_argument("-f", "--fxm-prefix", type=str, help="Prefix of the fxm file name (e.g. FITC, for FITC-1.tif file)")
parser.add_argument("-m", "--marker-prefix", type=str,
                    help="Prefix of the marker file name (e.g. mCherry, for mCherry-1.tif file)")

# Changes the following defaults to match your image filenames
parser.set_defaults(
    groups=2,
    fxm_prefix="GFP",
    marker_prefix="dsRED"
)

args = parser.parse_args()

path = args.path
global ng
ng = args.groups
fxm_prefix = args.fxm_prefix
marker_prefix = args.marker_prefix


if os.path.isdir(path):
    print("Run main.py first to calculate the volumes!")
    print("Then run group.py /path/to/data.tsv -g group_number")
    exit()

elif not os.path.isfile(path):
    print(f"Path '{path}' does not exist")
    exit()

print(f"Grouping cells in {ng} groups. Using '{fxm_prefix}*.tif' FXm images and '{marker_prefix}*.tif' marker images")

# Opens .tsv file
df = pd.read_csv(path, sep='\t', index_col=0)
# Extracting image filenames from analysis file
filenames = list(df["Path"].value_counts().index)
filenames.sort()

# Opens image by image in the interactive window
for file in filenames:
    img = mh.imread(file + ".tif")

    marker_file = file.replace(fxm_prefix, marker_prefix) + ".tif"
    marker = mh.imread(marker_file)

    mask_file = file + "_maskFram1.png"
    mask = mh.imread(mask_file)

    partial_df = df.loc[df['Path'] == file]

    groups = filter_cells(img, marker, mask, partial_df, path=file)

    df.loc[df['Path'] == file, "Group"] = groups


df["Group"] = df["Group"].astype(int)

counts = df["Group"].value_counts().sort_index()
print()
print("Group information:")
print(f"Total = {sum(counts)}")
print(counts)

# Saves .tsv with new column
df.to_csv(path.replace(".tsv", "_grp.tsv"), sep="\t")
```
