## Supplemental code for FXm analysis for "Fluorescence exclusion: a rapid, accurate and powerful method for measuring yeast cell volume": interactive.py

```
import numpy as np
import matplotlib.pyplot as plt
import autoSegment as auto
import mahotas as mh
import pandas as pd


def on_click(event):
    """
    Executed when user clicks on an object.
    Changes status of the object from selected to deselected (or vice versa)
    and draws a red line over the image.
    """
    global ix
    ix = event.xdata
    axes = event.inaxes

    if not axes or not axes in ax:  # Catches when user does not click on an image
        return

    # Finds clicked axes
    index = ax.index(axes)

    if preselection[index] is False:
        axes.plot(range(200), range(200), '-', linewidth=3, color="red")
        plt.draw()
        preselection[index] = True
    else:
        axes.lines[-1].remove()
        plt.draw()
        preselection[index] = False


def filter_cells(img, mask, cells_df, path=None):
    """
    Displays segmented objects. Allows for manual filtering
    of irrelevant objects based on their CALCULATED VOLUME

    :param img: original image
    :param mask: original segmentation mask
    :param cells_df: DataFrame with volume data
    :param path: path of the analyzed files. Will be used as the plot title
    :return cells_df: DataFrame with updated "ManualFilter" column (True for deselected objects)
    """

    img_size = img.shape[0]

    n_cells = len(cells_df["Volume"])

    # List of objects to be excluded. Updated upon user click.
    global preselection
    preselection = [False] * n_cells

    # Plotting parameters
    rows = np.ceil(np.sqrt(n_cells))
    columns = rows
    fig = plt.figure(figsize=(16, 9))  # Adapts these parameters for smaller screens
    fig.suptitle(path)

    centers = cells_df[["Center X", "Center Y"]].to_numpy()

    # Saves subplots in a list to be able to access them
    global ax
    ax = []

    j = 0
    for i, pix in enumerate(centers):

        # Adds new empty plot to selection window
        subp = fig.add_subplot(rows, columns, j + 1)

        # Formatting options
        subp.set_yticklabels([])
        subp.set_xticklabels([])
        subp.set_title(f"Cell: {str(i)} ({int(cells_df['Volume'][i])} µm3)")
        ax.append(subp)

        # Gets center coordinates of object
        x = int(pix[0])
        y = int(pix[1])

        # Gets 200x200 pixel area around object
        [x0, x1, y0, y1] = auto.get_bg_box(x, y, img_size=img_size)
        select = img[x0:x1, y0:y1]
        msk_img = select * (mask[x0:x1, y0:y1] > 0)

        plt.imshow(msk_img)

        # Uncomments next line to draw point to identify cell
        # plt.scatter(100, 100, s=1, color="red")

        j += 1  # Updates counter of displayed objects

    cid = fig.canvas.mpl_connect('button_press_event', on_click)  # Listens for click on object

    plt.show()

    cells_df["ManualFilter"] = preselection

    return cells_df
```
