## Supplemental code for FXm analysis for "Fluorescence exclusion: a rapid, accurate and powerful method for measuring yeast cell volume": main.py

```
import numpy as np
import pandas as pd
import os
import argparse

from scipy.io import loadmat
import autoSegment as auto
import interactive as inter


def analyze_experiment(path, manual=False, pillar_height=5.6, pixel_size=0.325):
    """
    Loads image and mask, calculates volumes and decides whether or not to analyze objects manually
    :param path: path to normalization file
    :param manual: manual analysis flag. If True, calls interactive functions to manually select the objects
    :param pillar_height: Height of the microfluidic chamber in µm
    :param pixel_size: Size of image pixel in µm given by your camera pixel size and the magnification used
    :return volumes: DataFrame with volume data and manual or automatic filter
    """

    # Load MATLAB normalization data (default)
    # If you have your own normalized images, load using mh.imread(os.path.join(path, "image.tif"))
    mat = loadmat(os.path.join(path, "frame1.mat"))
    image = mat["imageFlat"]
    mask = mat["deadZoneMask"]

    mask = mask > 0

    # Output variables
    segm_path = os.path.join(path, "../Segmentation")
    segm_file = "py_data_Auto.tsv"

    # Gets selections from normalization mask
    cells = auto.segment(mask)

    # Calculates volumes
    volumes = auto.get_volume(image, mask, cells, pillar_height=pillar_height, pixel_size=pixel_size)

    if volumes.empty:
        return pd.DataFrame({})

    if manual:
        # Displays objects for manual filtering by the user
        volumes = inter.filter_cells(image, mask, volumes, path=path)
        segm_file = f"py_data_Manual.tsv"

    # Adds folder name column
    volumes['Path'] = os.path.abspath(os.path.join(path, ".."))

    volumes.to_csv(os.path.join(segm_path, segm_file), sep="\t")  # Save

    return volumes


# User interaction and parameter logic
parser = argparse.ArgumentParser(description="Analyze images for S. pombe volume measurement.")
parser.add_argument("path", type=str, help="Path to images directory")
parser.add_argument("pillar", type=float, help="Height of the microfluidic chamber in µm")
parser.add_argument("--pixel", type=float, help="Size of image pixel in µm given by your camera pixel size and the "
                                                "magnification used", required=False)
parser.add_argument("-m", "--manual", action="store_true", help="Use manual filtering instead of automatic detection.")


# Changes defaults depending on your setup
thresholds = [1]  # List of IQR thresholds to use for automatic outlier detection

parser.set_defaults(
    pixel=0.325,  # Pixel size of your images (µm)
)

args = parser.parse_args()

analysis_dir = args.path
pillar_height = args.pillar
pixel_size = args.pixel
manual = args.manual

if not os.path.isdir(analysis_dir):
    print(f"The analysis directory does not exist.")
    exit()


norm_file = "frame1.mat"  # Normalization file that the script will look for


df = pd.DataFrame({})

# File finding
for root, dirs, files in os.walk(analysis_dir):
    for file in files:
        if file.endswith(norm_file):
            print(root)
            try:
                v = analyze_experiment(root, manual, pillar_height=pillar_height, pixel_size=pixel_size)
            except Exception as e:
                print(f"Image could not be analyzed due to an exception:\n{e}")
                continue
            if v.empty:
                print(f"No cells in image")
                continue
            if df.empty:
                df = v
            else:
                df = df.append(v, ignore_index=True)

print()

if df.empty:
    print(f"No data obtained. Did not find any {norm_file} files inside the analysis directory.")
    print(f"Have you normalized your images?")
    exit()

strain_dir = os.path.abspath(os.path.join(root, "../.."))
df_file = strain_dir + f"_A.tsv"

# Saves data and prints output information
if manual:
    df_file = strain_dir + f"_M.tsv"

    print("Manually filtered data:")
    print(df.loc[df["ManualFilter"] == False, "Volume"].describe())
    print()
else:
    df_file = strain_dir + f"_A.tsv"

    # Calculates IQR
    Q1 = np.percentile(df["Volume"], 25)
    med = np.percentile(df["Volume"], 50)
    Q3 = np.percentile(df["Volume"], 75)
    IQR = Q3 - Q1

    for th in thresholds:
        # Finds outliers
        outliers = np.logical_or(df["Volume"] > (Q3 + IQR * th), df["Volume"] < (Q1 - IQR * th))
        filter_name = f"AutoFilterIQR_{th}"  # Name of the column that includes IQR size
        df[filter_name] = outliers
        print(f"Automatically filtered data (IQR {th}):")
        print(df.loc[df[filter_name] == False, "Volume"].describe())
        print()

print(f'Output file: {df_file}')
df.to_csv(df_file, sep="\t")
```
