## Supplemental code for FXm analysis for "Fluorescence exclusion: a rapid, accurate and powerful method for measuring yeast cell volume": README.md

# select-pombe-fxm

This code facilitates the analysis of data obtained using the Fluorescence Exclusion Method (FXm) for the measurement of yeast cell volume. 


## Dependencies

This code requires the following Python packages (also see `requirements.txt`):
```
mahotas==1.4.9
matplotlib==3.2.1
numpy==1.18.5
pandas==1.0.4
scipy==1.6.0
```

## How does it work?

FXm analysis involves a step of image normalization.
This step generates a mask in which objects are pre-identified.

select-pombe-fxm uses this mask and:

1. Automatically selects all objects and filters out those that show an abnormally high or low volume, or

2. Displays all pre-selected objects to the user for manual filtering

3. Optional: assigns different groups to the selected cells, based on other fluorescent markers


The normalization mask is generated using a custom MATLAB software from Cadart et al., Methods Cell Biol. (2017).


## Installation

1. Create a virtual environment and activate it.
    ```
    python3.7 -m venv venv
    source venv/bin/activate
    ```
2. Install the requirements in `requirements.txt`.
    ```
    pip install -r requirements.txt
    ```


## Usage
```shell script
# Activate virtual environment
source venv/bin/activate
```
```shell script
# Run script to calculate volumes
python main.py -h
usage: main.py [-h] [--pixel PIXEL] [-t [THRESHOLDS] | -m] path pillar

Analyze images for S. pombe volume measurement

positional arguments:
  path                  Path to images directory
  pillar                Height of the microfluidic chamber in µm

optional arguments:
  -h, --help            Shows this help message and exit
  --pixel PIXEL         Size of image pixel in µm given by your camera pixel
                        size and the magnification used
  -m, --manual          Uses manual filtering instead of automatic detection

```
```shell script
# Optional: run script to assign groups to cells
python group.py -h
usage: group.py [-h] [-g GROUPS] [-f FXM_PREFIX] [-m MARKER_PREFIX] path

Separate FXm cells in groups

positional arguments:
  path                  Path to .tsv analysis file

optional arguments:
  -h, --help            Shows this help message and exit
  -g GROUPS, --groups GROUPS
                        Number of groups to classify the cells in
  -f FXM_PREFIX, --fxm-prefix FXM_PREFIX
                        Prefix of the fxm file name (e.g. FITC, for FITC-1.tif file)
  -m MARKER_PREFIX, --marker-prefix MARKER_PREFIX
                        Prefix of the marker file name (e.g. mCherry, for mCherry-1.tif file)

```

### Automatic mode

This mode automatically selects all objects and discards those that show abnormally high or low volumes (1 IQR, corresponds to 2 SD threshold)

Enter the path and pillar height arguments and run:
```
python main.py </path/to/experiment/files> <pillar-height>
```

This will generate a `.tsv` file with all the data.
This file includes a `bool` column with the results of the automatic filtering.

`True`: outlier - data discarded

`False`: valid cells


### Manual mode

This mode allows the user to manually exclude irrelevant objects from the analysis.

Run the manual mode by adding the `-m` flag:
```
python main.py </path/to/experiment/files> <pillar-height> -m
```

This will open a `pyplot` window showing all pre-selected objects of the image.
Irrelevant objects can be de-selected by simple mouse click (discarded objects are red-barred).
When the manual filtering of a given image is done, simply close the window (data are automatically saved).
A new window for the next image is then opened.

Note: each subplot is a 200x200 px image centered on the object that is being selected.
However, these selections do not need to be discarded as only the centered object will be considered for volume calculation.

When all images have been treated, a `.tsv` file containing all data is generated.
It includes a column with the results of the manual filtering.
`True`: manually discarded as outlier
`False`: valid cell

### Group mode
This mode allows the user to separate cells into different groups, based on other fluorescent markers.
For this, it uses 

Run the group mode by:
```shell script
# To assign groups to manually filtered cells (run Manual mode first)
python group.py </path/to/data_M.tsv> -g <group number> -f <prefix of FXm images> -m <prefix of marker images>
```
```shell script
# To assign groups to all cells (run Automatic mode first)
python group.py </path/to/data_A.tsv> -g <group number> -f <prefix of FXm images> -m <prefix of marker images>
```
This will open a `pyplot` window showing the selected cells or all the objects, which are automatically assigned to Group 1 (G1).
The user will be able to select the appropriate group as follows:

- Left click on each object to increase the group number.
- Right click on each object to decrease the group number.
- Discard object by assigning the `None` group (group 0) (for instance, right click when group is 1)
- Click on the radiobuttons (or press `space`) to alternate between FXm image and the fluorescent marker image.
